# Population Structure Discovery in Meta-Analyzed Microbial Communities and Inflammatory Bowel Disease

**DOI:** 10.1101/2020.08.31.261214

**Authors:** Siyuan Ma, Dmitry Shungin, Himel Mallick, Melanie Schirmer, Long H. Nguyen, Raivo Kolde, Eric Franzosa, Hera Vlamakis, Ramnik Xavier, Curtis Huttenhower

## Abstract

Microbial community studies in general, and of the human microbiome in inflammatory bowel disease (IBD) in particular, have now achieved a scale at which it is practical to associate features of the microbiome with environmental exposures and health outcomes across multiple large-scale populations. This permits the development of rigorous meta-analysis methods, of particular importance in IBD as a means by which the heterogeneity of disease etiology and treatment response might be explained. We have thus developed MMUPHin (Meta-analysis Methods with a Uniform Pipeline for Heterogeneity in microbiome studies) for joint normalization, meta-analysis, and population structure discovery using microbial community taxonomic and functional profiles. Applying this method to ten IBD cohorts (5,151 total samples), we identified a single consistent axis of microbial associations among studies, including newly associated taxa such as *Acinetobacter* and *Turicibacter* detected due to the sensitivity of meta-analysis. Linear random effects models further revealed associations with medications, disease location, and interaction effects consistent within and between studies. Finally, multiple unsupervised clustering metrics and dissimilarity measures agreed on a lack of discrete microbiome “types” in the IBD gut microbiome. These results thus provide a benchmark for consistent characterization of the IBD gut microbiome and a general framework applicable to meta-analysis of any microbial community types.

## Introduction

Meta-analysis for molecular epidemiology in large populations has seen great success in linking high-dimensional ‘omic features to complex health-related phenotypes. One example of this is in genome-wide association studies (GWAS^1^), where the appropriate study scale, achieved by rigorous integration of multiple cohorts, has both facilitated reproducible discoveries (in the form of disease-associated loci^2-4^) and addressed confounding due to unobserved population structure^5^. The inflammatory bowel diseases (IBD) represent a particular success story for GWAS meta-analysis^3,4^, and environmental and microbial contributors complementing the condition’s complex genetic architecture have been detailed by many individual studies^6-8^. However, in the absence of methods appropriate for large-scale microbial meta-analysis, the extent to which these findings reproduce across studies, or can be extended by increased joint sample sizes, remains undetermined. Likewise, it is unclear whether reproducible population structure in the microbiome, such as microbially-driven IBD “subtypes,” exists to help explain the clinical heterogeneity of these conditions^9^.

Meta-analysis of microbial community profiles presents unique quantitative challenges relative to other types of ‘omics data such as GWAS^10^ or gene expression^11^. These include particularly strong batch, inter-individual, and inter-population differences, and statistical issues including zero-inflation and compositionality^12,13^. Consequently, methods to correct for cohort and batch effects from other ‘omics settings^14-17^ are not directly appropriate. Two recent studies have suggested quantile normalization^18^ and Bayesian Dirichlet-multinomial regression (BDMMA)^19^ for microbial profiles, which are applicable to a limited subset of differential abundance tests and do not provide batch-corrected profiles. To date, there are no methods permitting the joint analysis of batch-corrected microbial profiles for most study designs.

IBD represents one of the best-studied, microbiome-linked inflammatory phenotypes to date which thus stands to benefit from such approaches^20,21^. Among the inflammatory bowel diseases, Crohn’s disease (CD) and ulcerative colitis (UC) have been individually linked with structural and functional changes in the gut microbiome in many individual studies^21^. Each of CD and UC can itself be highly heterogeneous within the IBD population, however, and diversity in disease-associated gut microbial features has not been consistently associated with factors including disease subtype, progression, or treatment response^7,9,22,23^. Of note, two meta-analysis studies included IBD as one of several phenotypes^24,25^. These studies were not IBD-specific, did not have access to appropriate normalization techniques, nor took the aforementioned factors into account. The complexity of microbial involvement in IBD, and the presence of substantial unexplained variation in the manifestation of its symptoms, makes it particularly appropriate for application of meta-analysis techniques.

In this work, we introduce and validate a statistical framework for population-scale meta-analysis of microbiome data, and apply it to the largest collection to date of ten published 16S rRNA gene sequencing-based IBD studies (**Table 1**) to identify consistent disease associations and population structure. We found both previously documented and novel microbial links to the disease, with further differentiation among subtypes, phenotypic severity, and treatment effects. We further confidently conclude that there are no apparent, reproducible microbiome-based subtypes within CD or UC, which are instead a population structure gradient from less to more “pro-inflammatory” ecological configurations. Our work thus represents one of the first large-scale efforts to assesses consistency in gut microbial findings for IBD and provides methodology supporting future microbial community meta-analyses.

**Table 1:** 10 uniformly processed 16S rRNA gene sequencing studies of the IBD mucosal/stool microbiomes. For longitudinal cohorts, numbers in parentheses indicate baseline sample size. For age, mean and standard error (parenthesized) are shown. Additional covariates are summarized in **Supplemental Table 1**.

| Study | Brief description | N subject | N sample | Phenotype(s) | Age | Gender | Sample type(s) |
| --- | --- | --- | --- | --- | --- | --- | --- |
| PROTECT<sup>23</sup> | Longitudinal cohort of newly diagnosed UC | 405 | 1212 (539) | UC 405 | 12.71 (3.29) | Male 52%/<br>Female 48% | Biopsy 22%/<br>Stool 78% |
| RISK<sup>7</sup> | Pediatric cohort of treatment-naïve CD | 631 | 882 | CD 430/<br>Control 201 | 12.16<br>(3.22) | Male<br>59%/<br>Female<br>41% | Biopsy<br>72%/<br>Stool<br>28% |
| Herfarth<sup>27</sup> | Densely (daily) sampled longitudinal cohort | 31 | 860<br>(31) | CD 19/<br>Control 12 | 36.03<br>(14.12) | Male<br>35%/<br>Female<br>58%/<br>Missing<br>6% | Stool |
| Jansson-Lamendella<sup>22</sup> | Longitudinal follow up with fecal samples | 137 | 683<br>(137) | CD 49/<br>UC 60/<br>Control 28 |  | Male<br>42%/<br>Female<br>58% | Stool |
| Pouchitis<sup>28</sup> | Patients recruited underwent IPAA for treatment of UC or FAP prior to enrollment. | 353 | 577 | CD 42/<br>UC 266/<br>Control 45 | 46.19<br>(13.58) | Male<br>52%/<br>Female<br>48% | Biopsy |
| CS-PRISM<sup>29</sup> | Cross sectional cohort nested in PRISM | 397 | 467 | CD 215/<br>UC 144/<br>Control 38 | 41.68<br>(15.22) | Male<br>47%/<br>Female<br>53% | Biopsy<br>29%/<br>Stool<br>71% |
| HMP2<sup>9</sup> | Large cohort of newly diagnosed IBD with multi 'omics measurement. | 81 | 177<br>(162) | CD 37/<br>UC 22/<br>Control 22 | 29.76<br>(19.63) | Male<br>51%/<br>Female<br>49% | Biopsy |
| MucosalIBD<sup>30</sup> | Pediatric cohort with Paneth cell phenotypes | 83 | 132 | CD 36/<br>Control 47 | 12.93<br>(3.65) | Male<br>58%/<br>Female<br>42% | Biopsy |
| LSS-PRISM<sup>31</sup> | Longitudinal cohort nested in PRISM. | 18 | 88 (19) | CD 12/<br>UC 6 | 30.37<br>(10.52) | Male<br>39%/<br>Female<br>61% | Stool |
| BIDMC-FMT<sup>32</sup> | FMT Trial design | 8 | 16 | CD 8 | 38.38<br>(12.73) | Male<br>62%/<br>Female<br>38% | Stool |

## Results

### Integrating 10 studies of the IBD stool and mucosal microbiomes

We collected and uniformly processed ten published 16S studies of the IBD gut microbiome (**Table 1, Fig. 1a, Supplemental Table 1**) totaling 2,179 subjects and 5,151 samples. These studies range widely in terms of cohort designs and population characteristics, including recent-onset and established disease patients, cross-sectional and longitudinal sampling, pediatric and adult populations, diseases (CD and UC), treated and treatment-naive patients, biopsy and stool samples, and inclusion of healthy/non-IBD controls. Covariates were manually curated to ensure consistency across studies (**Methods**). Major factors available from all or most studies included demographics (age/sex/race), biogeography, disease location and/or extent, antibiotic usage, immunosuppression, and steroid and/or 5-ASA usage.

**Figure 1:**
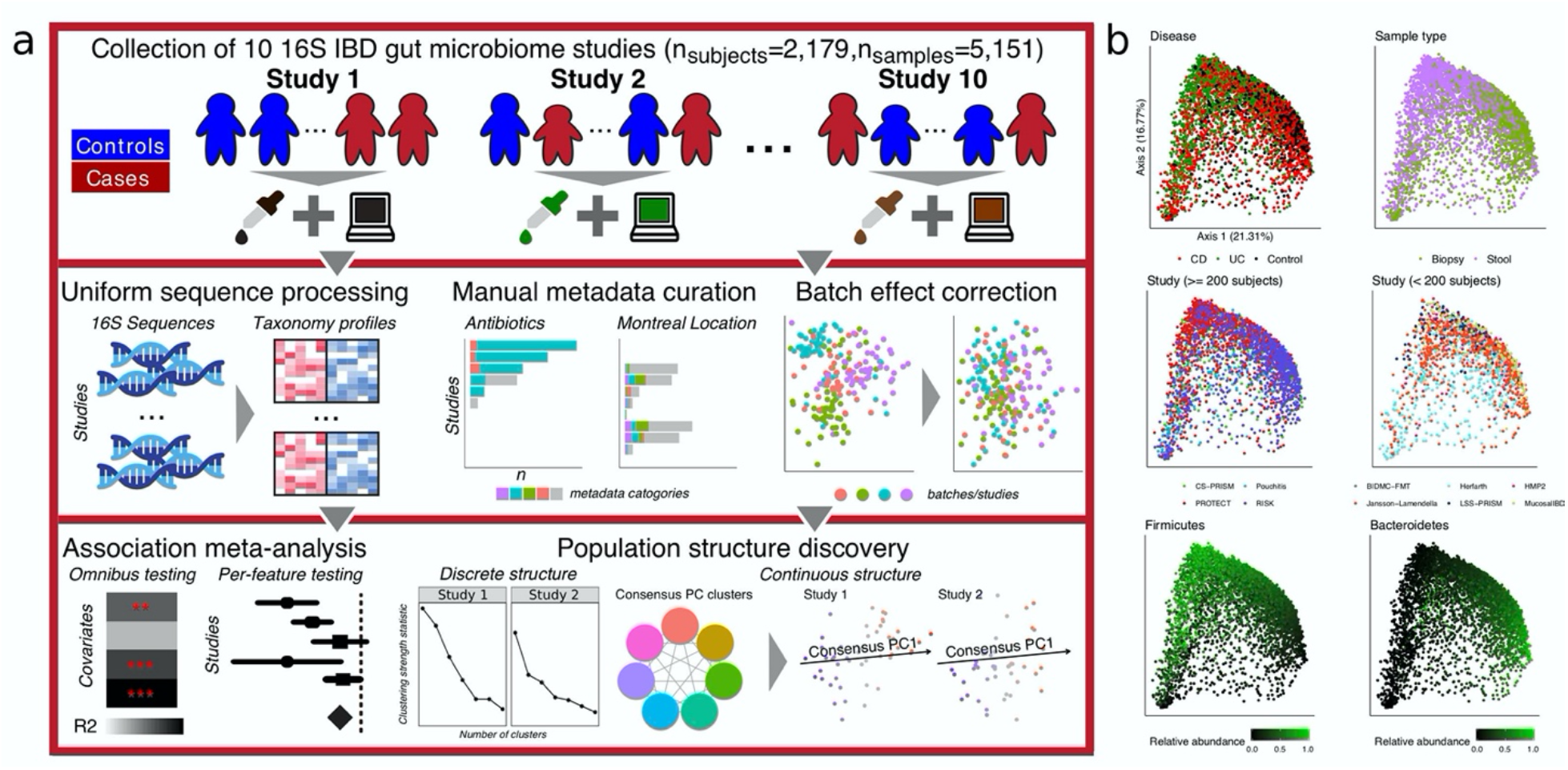
A method for large-scale microbial community meta-analysis and its application to inflammatory bowel disease. **a)** We developed a novel statistical framework, MMUPHin, allowing joint normalization and meta-analysis of large microbial community profile collections with heterogeneous and complex designs (multiple covariates, longitudinal samples, etc.). We applied it to a collection of 10 inflammatory bowel disease studies comprising 2,179 subjects and 5,151 total samples (**Table 1**). We uniformly processed the associated sequence data and harmonized metadata across cohorts. Microbial taxonomic profiles were then corrected for batch- and study-effects before downstream analyses for omnibus and per-feature association with disease phenotypes and unsupervised population structure discovery. **b)** MDS ordination of all microbial profiles (Bray-Curtis dissimilarity) before batch correction visualize the strongest associations with gut microbial composition, including disease, sample type (biopsy or stool), cohort (visualized separately for larger and smaller studies), and dominant phyla.

Using this joint dataset and upon uniform bioinformatics processing (**Methods**), we first assessed the factors that corresponded to overall variation in microbiome structure, which included disease status, sample type (biopsy versus stool), and dominant phyla (Bacteroidetes and Firmicutes, **Fig. 1b**). Cohort effects prior to batch correction and meta-analysis were also significant. Microbiome differences associated with disease were notable even without normalization. However, this can be misleading due to the confounding of cohort structure between studies, such as the differentiation between RISK (a predominantly mucosal study of CD) and PROTECT (a predominantly stool study of UC). Inter-individual differences largely independent of population or disease, such as Bacteroidetes versus Firmicutes dominance, were also universal among studies and sample types as expected^9,26^. Many of these factors were of comparable effect size, both visually and as quantified below, emphasizing the need for covariate-adjusted statistical modelling to delineate the biological (disease, treatment) and technical (cohort, batch) effects associated with individual taxa throughout the cohorts (**Supplemental Notes, Supplemental Fig. 1-3**).

### A statistical framework for meta-analysis of microbial community profiles

We developed a collection of novel methods for meta-analysis of environmental exposures, phenotypes, and population structures across microbial community studies, specifically accounting for technical batch effects and interstudy differences (**Methods, Fig. 1a**). It consists of three main components: batch and study effect correction, covariate modeling, and population structure discovery. First, we extended methods from the gene expression literature (ComBat^15^) to enable batch correction of zero-inflated microbial abundance data. Based on linear modelling, the method can differentiate between technical effects (batch, study) versus covariates of biologically interest (exposure, phenotype). Second, we combined well-validated data transformation and linear modelling combinations for microbial community profiles^33^ with fixed and random effect modelling^34^ for meta-analytical synthesis of per-feature (taxon, gene, or pathway) differential abundance effects. Lastly, we generalized and formalized approaches from cancer transcriptional subtyping^35^ to permit unsupervised discovery and validation of both discrete and continuous population structures in microbial community data (**Supplemental Fig. 4**). Our methods, implemented as Meta-analysis Methods with a Uniform Pipeline for Heterogeneity in microbiome studies (MMUPHin), are available as an R package through Bioconductor^36^ and at https://bioconductor.org/packages/release/bioc/html/MMUPHin.html.

We validated MMUPHin both in comparison to existing methods and through extensive simulation studies (**Fig. 2**), with simulated realistic microbial abundance profiles at different data dimensionality, biological/technical batch signal strength, and discrete/continuous population structures (**Methods, Supplemental Table 2, Supplemental Fig. 5-8**). MMUPHin successfully reduced variability attributable to technical effects in simulated microbial profiles, as first quantified by the PERMANOVA R2 statistic^37^ (**Fig. 2a-b, Supplemental Fig. 5**). This was true both in terms of reducing the overall microbial variability attributable to technical artifacts and in terms of the ratio of “biological” versus technical variability (**Fig. 2a**). ComBat correction^15^, suited for gene expression data, was capable of reducing batch effects to a lesser degree, but also tended to reduce desirable “biological” variation in the process, likely due to noise introduced by it changing many zero counts to non-zero values. Previously proposed techniques for microbial community data, namely quantile normalization^18^ and BDMMA^19^, are only appropriate for differential abundance analysis and do not provide batch-normalized profiles, thus precluding PERMANOVA batch effect quantification; their per-feature testing performance is evaluated together with MMUPHin in the following section. MMUPHin thus provides batch-corrected microbial community profiles that retain biologically meaningful variation more than (or not even possible using) existing methods.

**Figure 2:**
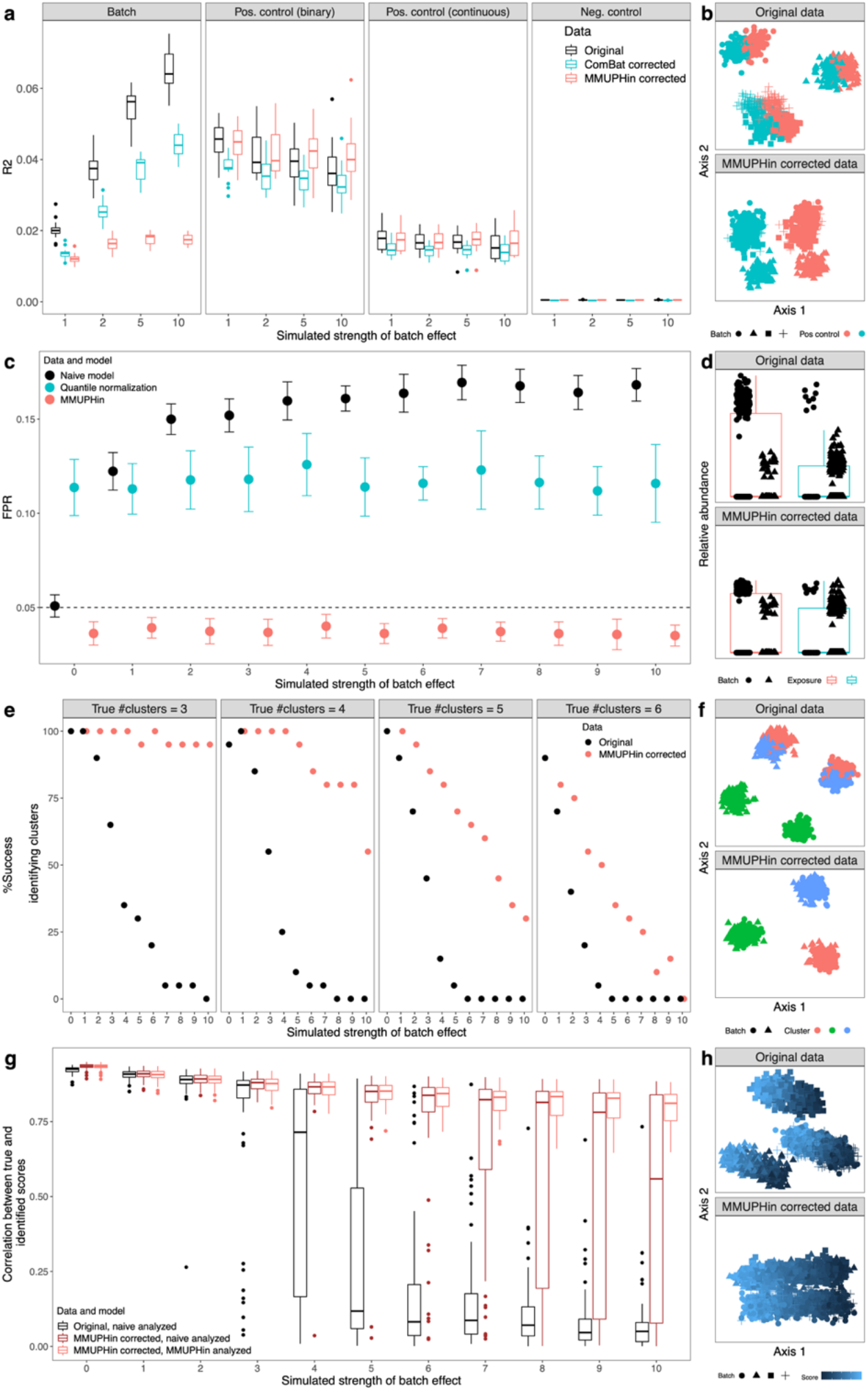
Effectiveness of batch correction, association meta-analysis, and unsupervised population structure discovery methods. All evaluations use simulated microbial community profiles as detailed in Methods. Left panels summarize representative subsets of results (full set of simulation cases presented in **Supplemental Table 2** and results in **Supplemental Fig. 5-8**), and right panels show examples of batch-influenced data pre- and post-correction. a, b) MMUPHin is effective for covariate-adjusted batch effect reduction while maintaining the effect of positive control variables. Results shown correspond to the subset of details in **Supplemental Fig. 5** with number of samples per batch = 500, number of batches = 4, and number of features = 1000 with 5% spiked with associations. c, d) Batch correction and meta-analysis reduces false positives when an exposure is spuriously associated with microbiome features due to an imbalanced distribution between batches. Corresponds to **Supplemental Fig. 6** with number of samples per batch = 500, number of features = 1000 with 5% spiked associations, and case proportion difference between batches = 0.8. Evaluations of BDMMA generates low FPRs due to the zero-inflated nature of simulated microbial abundances, and are included only in **Supplemental Fig. 6**. e, f) Batch correction improves correct identification of the true underlying number of clusters during discrete population structure discovery. Corresponds to **Supplemental Fig. 7** with number of batches = 4. g, h) Continuous structure discovery accurately recovers microbiome compositional gradients in a simulated population. Corresponds to **Supplemental Fig. 8** with number of batches = 6.

For differential abundance testing, MMUPHin successfully corrected for false associations when batch/cohort effect<u>s</u> were confounded with variables of interest, which is a common concern for ‘omics meta-analysis^38^, while quantile normalization^18^ and BDMMA^19^ had either inflated or overly conservative false positive rates (**Fig. 2c-d, Supplemental Fig. 6**). We also validated MMUPHin’s support for unsupervised population structure discovery, in addition to these “supervised” differential abundance and statistical association tests. In microbial communities, valid, generalizable population structure can manifest as either discretely clustered subtypes^39^ or as continuously variable gradients of community configurations^40^, but methods for discovery are particularly susceptible to false positives in the presence of technical artifacts^26,40^. To this end, for discrete structures, MMUPHin utilizes established clustering strength evaluation metrics^41^ to a) evaluate the existence of discrete clusters within individual microbiome studies and b) to validate the reproducibility of such structures among studies meta-analytically (**Fig. 2e-f, Supplemental Fig. 7**). For continuous structures, our method generalizes single study principal component analysis (PCA^42^) to multiple studies by constructing a network of correlated top PC loadings^35^, thus identifying major axes of variation that explain the largest amount of heterogeneity between microbial profiles and are also consistent across studies (**Fig. 2g-h, Supplemental Fig. 8**). As a result, MMUPHin was able to successfully identify discrete clusters (i.e. microbiome “types”) when present, as well as significantly consistent continuous patterns of microbiome variation that recur among populations (**Supplemental Notes**).

### Meta-analysis of the IBD microbiome

Given these validations of MMUPHin’s accuracy in simulated data, we next applied it to the 10-study, 4,789-sample IBD gut amplicon profile meta-analysis introduced above (**Fig. 3**). MMUPHin successfully reduced the effects both of differences among studies, and of batches within studies (study effect correction modelling disease and sample type as covariates, see **Methods**), although these remained among the strongest source of variation among taxonomic profiles as quantified by PERMANOVA R2 (**Fig. 3a, Methods, Supplemental Table 3**). Among biological variables, sample type (biopsy/stool), biopsy location (multiple, conditional on biopsy samples), disease status (IBD/control), and disease types (CD/UC, conditional on IBD) consistently had the strongest effect on the microbiome among studies. Several relationships between study design and phenotypic effects were apparent. Batches had a particularly strong effect in CS-PRISM and RISK, for example, where biopsy and stool samples were also perfectly separated by batch.

**Figure 3:**
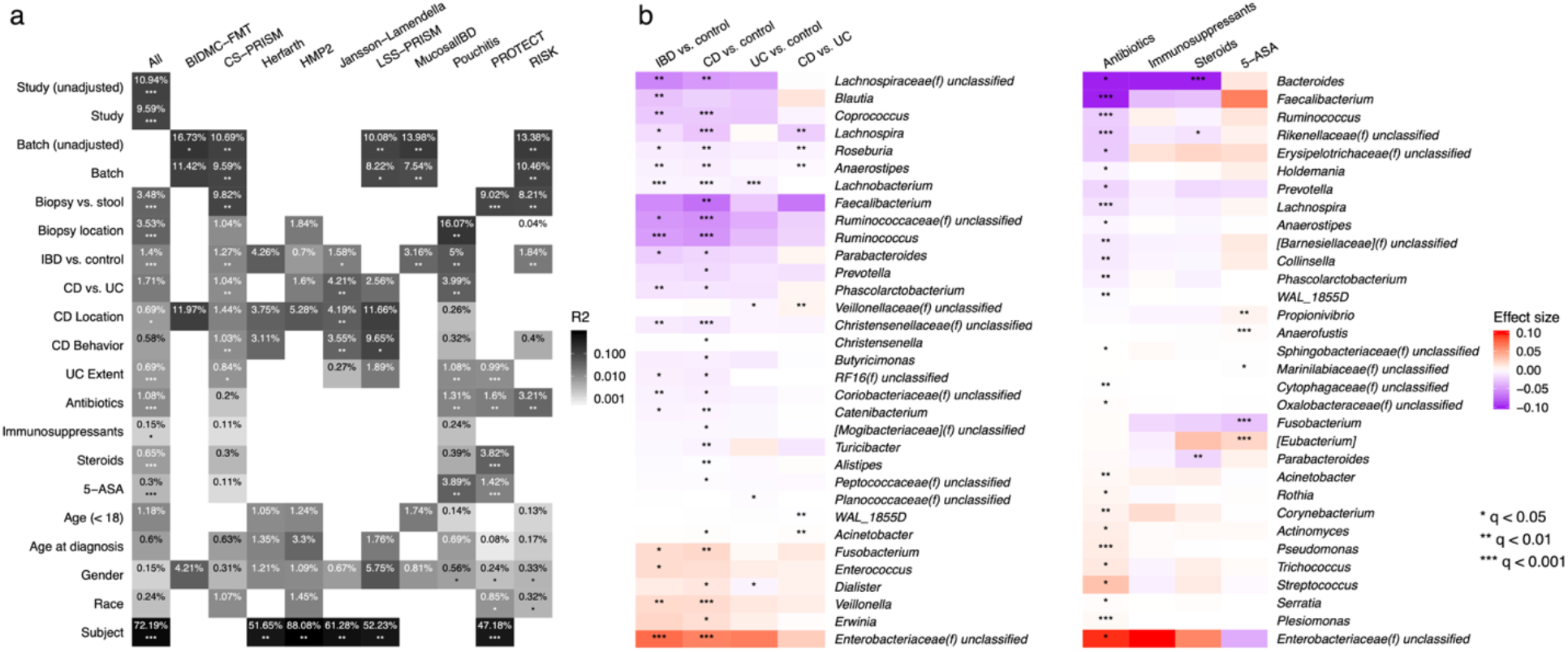
Meta-analytic omnibus and per-feature testing reveal novel and previously documented IBD associations. **a)** Omnibus testing (PERMANOVA on Bray-Curtis dissimilarities with stratification and covariate control where appropriate, see **Methods** and **Supplemental Table 3**) identified between-subject differences as the greatest source of microbiome variability, with IBD phenotype, disease (CD/UC), and sample type (stool/biopsy) as additional main sources of biological variation. MMUPHin successfully reduced between-cohort and within-study batch effects, although these technical sources also remained significant contributors to variability. **b)** Individual taxa significantly associated with IBD phenotypes or treatments after meta-analysis. Taxa are arranged by family-level median effect size of IBD vs. control for disease results and that of antibiotic usage for treatment results. Effect sizes are aggregated regression coefficients (across studies with random effects modelling) on arcsin square root-transformed relative abundances. Detailed model information in **Methods** and **Supplemental Table 3**. Individual study results in **Supplemental Table 4**.

Treatment exposures all had small effects on microbiome structure within studies, which typically reached statistical significance only when combined by meta-analysis; antibiotics were an exception with slightly larger effects. Montreal classification did not generally correspond with significant variation, while age (at sample collection as stratified below and above 18, and at diagnosis by Montreal age classification^43^) had small but significant effects. The effects of gender and race were not significant. Lastly, for longitudinal studies, relatively stable differences between subjects over time were large and significant, consistently for both longer-interval (HMP2) as well as densely sampled cohorts (Herfarth, daily samples), in agreement with previous individual studies’ observations^9,23^.

We identified individual taxonomic features consistently associated with disease and treatment variables (**Fig. 3b, Supplemental Table 4**), with meta-analysis multivariate differential abundance analysis, adjusting for common demographics (age, gender, race) and further stratifying for sample type and disease when appropriate (**Methods, Supplemental Table 3**). At a very high level, differential abundance patterns between CD and control microbiomes were consistent with, and often more severe than contrasts between UC and control, confirming with increased resolution previous observations that CD patients tend to have more aggravated dysbiosis than UC patients^9^. As expected, our meta-analysis confirms many of the taxa associated with IBD reported by previous individual (**Fig. 3b**, detailed in **Supplemental Notes**); these findings strongly supports the emerging hypotheses of pro-inflammatory aerotolerant clades forming a positive feedback loop in the gut during inflammation, often of oral origin^7^, and depleting the gut’s typical fastidious anaerobe population as a result.

We also identified two taxa not previously associated with IBD, both of modest effect sizes and likely newly detected by the meta-analysis’ increased power. The genus *Acinetobacter* was enriched in CD, and *Turicibacter* was depleted. *Turicibater* in particular is poorly represented in reference sequence databases, with only nine genomes for one species (*Turicibacter sanguinis*) currently in the NCBI genome database; this makes it easy to overlook in shotgun metagenomic profiles relative to amplicon sequencing. The genus *Acinetobacter*, conversely, is quite well characterized due to its role in antimicrobial resistant infections^44^, and it was previously linked specifically to the primary sclerosing cholangitis phenotype in UC^45^, although without follow-up to our knowledge. *Turicibacter* is overall less characterized both in isolation and with respect to disease, although our findings and others’ suggest it might be inflammation-sensitive when present; it was one of many clades increased in mice during CD8+ T cell depletion^46^ and reduced in a homozygous TNF deletion^47^. As the strains of *Acinetobacter* implicated in gut inflammation are unlikely to be those responsible for e.g. nosocomial infections, further investigation of both clades using more detailed data or IBD-specific isolates is warranted.

Among treatment variables (samples or time points during which subjects were receiving antibiotics, immunosuppressants, steroids, and/or 5-ASAs), antibiotics had the strongest effects on individual taxa, as well as the greatest number of significantly associated taxa (**Fig. 3b**). These associations are also broadly in agreement with previous observations for microbiome responses to antibiotics in IBD or generally, e.g. the depletion of *Faecalibacterium, Ruminococcus*, and *Bacteroides* in patients treated with antibiotics, and the enrichment of (often stereotypically resistant) taxa such as *Streptococcus, Acinetobacter*, and the Enterobacteriaceae, with differential responses to the treatment groups speaking to both administration considerations and their impact on host versus microbial community bioactivities (**Supplemental Notes**).

Subsets of IBD-linked taxa were additionally associated with the diseases’ phenotypic severity (**Fig. 4a, Supplemental Table 5**). Montreal classification^43^ was used as a proxy for disease severity, including Behavior categories for Crohn’s disease (B1 non-stricturing, non-penetrating, B2 stricturing, non-penetrating, B3 stricturing and penetrating) and Extent for ulcerative colitis (E1 limited to rectum, E2 up to descending colon, E3 pancolitis). We tested for features differentially abundant in the more severe phenotypes when compared against the least severe category (B1 CD and E1 UC, **Methods**). Among statistically significant results, many extended those identified above as overall IBD associated (**Fig. 3b**), such as the depletion of *Faecalibacterium* in B3 CD and *Roseburia* in B2 CD, as well as the enrichment of Enterobacteriaceae in E3 UC. In most cases, microbial dysbiosis was also additionally aggravated from the moderate to the most extreme disease manifestations; such differences were statistically significant (**Methods**) in, for example, the progressive depletion of *Bacteroides* in CD and UC, as well as the enrichment of Enterobacteriaceae in UC. This meta-analysis is uniquely powered to detect these subtle differences, which aid in shedding light on the microbiome’s response to progressive inflammation and disease subtypes. Pancolitis corresponds with a unique microbial configuration distinct from regional colitis and not generally detectable in smaller studies^6^, for example, while more severe CD induces essentially a more extreme form of the same dysbiosis observed in less severe forms of the disease.

**Figure 4:**
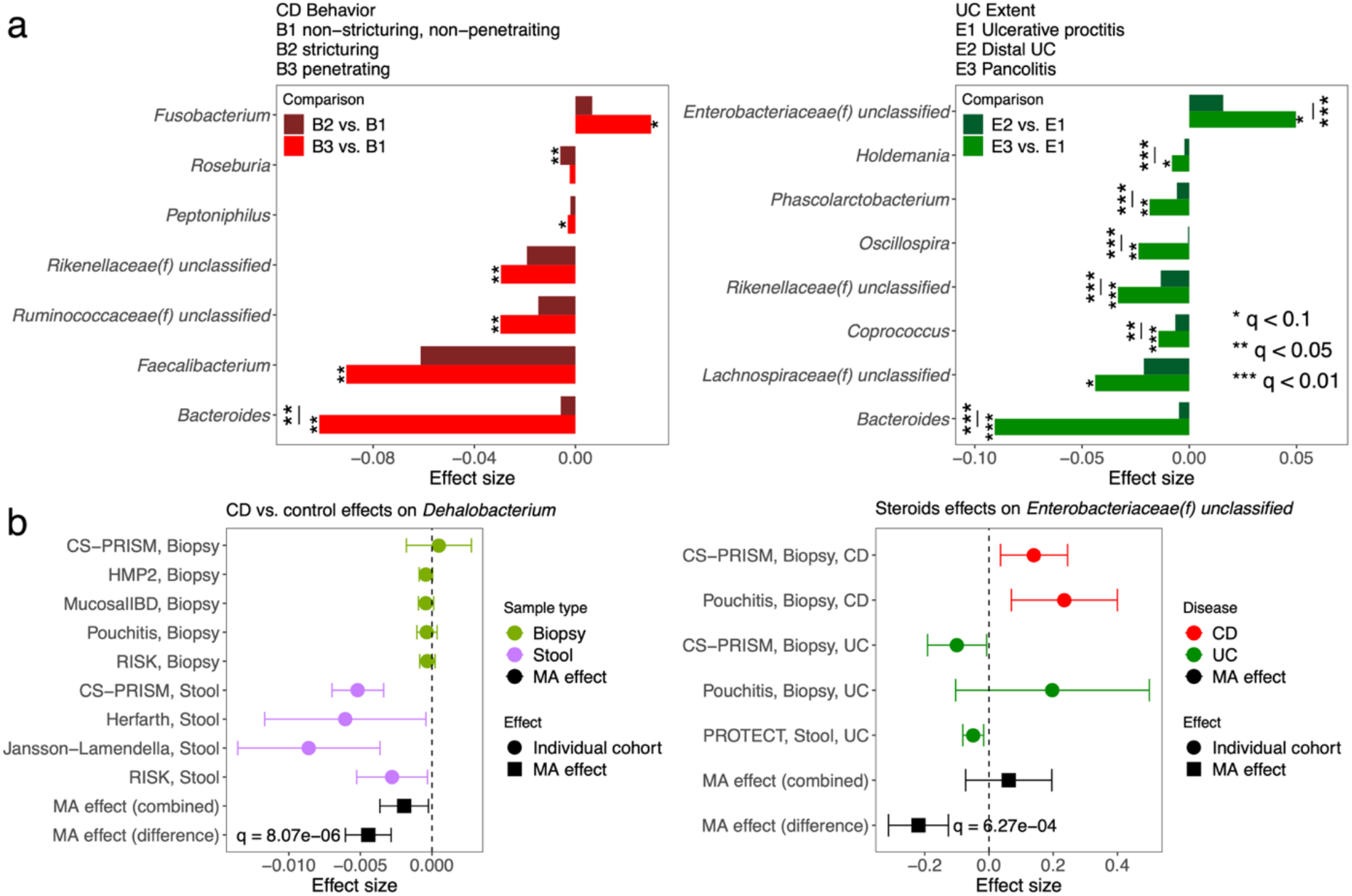
IBD-associated taxa are aggravated in more severe disease; disease biogeography and CD/UC differentially affect some taxa with respect to disease and treatment. **a)** Statistically significant genera from meta-analytically synthesized differential abundance effects among severity of CD and UC phenotypes as quantified by Montreal classification. The difference between the most severe phenotype with the least severe one (B3 vs. B1 for CD, E3 vs. E1 for UC) was in most cases more aggravated than that of the intermediate phenotype. Many of the identified features overlap with those associated with IBD vs. control differences, suggesting a consistent gradient of severity effects on the microbiome. Individual study results in **Supplemental Table 5. b)** Genus *Dehalobacterium* as an example in which a taxon is uniquely affected in the stool microbiome during CD and not at the mucosa. Likewise, family Enterobacteriaceae as an example in which steroid treatment corresponds with enrichment of the clade in CD samples, but depletion in UC. In all panels, effect sizes are aggregated regression coefficients on arcsin square root-transformed relative abundances. Full sets of statistically significant interactions, with individual study results, are in **Supplemental Table 6**.

Additionally, diseases (CD and UC) and their corresponding dysbioses also interacted distinctly with the microbiome under different treatment regimes and in different biogeographical environments (mucosa vs. stool, **Fig. 4b, Supplemental Table 6**). Interaction effects, in the statistical sense, were defined as a main exposure (IBD or treatment) having differential effects on taxon abundance with respect to either sample type (biopsy/stool) or diseases (CD/UC); they were identified via moderator meta-analysis models (**Methods**). Overall, we found elevated effects of both CD (relative to controls) and antibiotic treatment in stool as compared to biopsy-based measurements of the microbiome (**Supplemental Table 6**). An example of this is *Dehalobacterium*, with significantly greater depletion in CD stool relative to biopsies (**Fig. 4b**). *Dehalobacterium*, as with *Turicibacter* above, is underrepresented in reference sequence databases, better-detected by amplicon sequencing, and thus not a common microbial signature of IBD. It has been linked to CD in at least one existing 16S-based stool study^48^. In contrast, several UC-specific microbial disruptions were more prominent at the mucosa (i.e. in biopsies, **Supplemental Table 6**). Coupled with the severity-linked differences above, this suggests CD-induced changes in the entire gut microbial ecosystem largely as a consequence of inflammation, with UC-induced dysbioses both more local and more specific to disease and treatment regime. Additional results include effect of steroids on the Enterobacteriaceae, which tended to be more abundant in CD patients receiving steroids, but less abundant in UC recipients (**Fig. 4b, Supplemental Table 6, Supplemental Notes**).

### Consistent IBD microbial population structure discovered by unsupervised analysis

The existence of subtypes within gut microbial communities has been a major open question in human microbiome studies, and it is of particular importance within IBD as a potential explanation for heterogeneity in disease etiology and treatment response^6,9^. To systematically characterize population structure in the IBD gut microbiome that was reproducible among studies, we performed both discrete and continuous structure discovery on the 10 cohorts using our meta-analysis framework. To identify potential discrete community types (i.e. clusters), we performed clustering analysis within each cohort’s IBD patient population, and evaluated the clustering strength via prediction strength (**Methods**). We found no evidence to support discrete clustering structure within individual cohorts, nor were we able to reproduce each cohort’s clustering results externally (**Fig. 5a**). This lack of discrete structure was consistent when we further stratified samples to either CD or UC populations (**Supplemental Fig. 9**), or extended to additional dissimilarity metric and clustering strength measurements (**Supplemental Fig. 9, Methods**). Our observation that the IBD gut microbiome cannot be well characterized by discrete clusters is thus consistent with previous findings on gut microbial heterogeneity for healthy populations^40^ and suggests that, at the level powered by this study, such microbiome subtypes are not clearly responsible for clinical heterogeneity.

**Figure 5:**
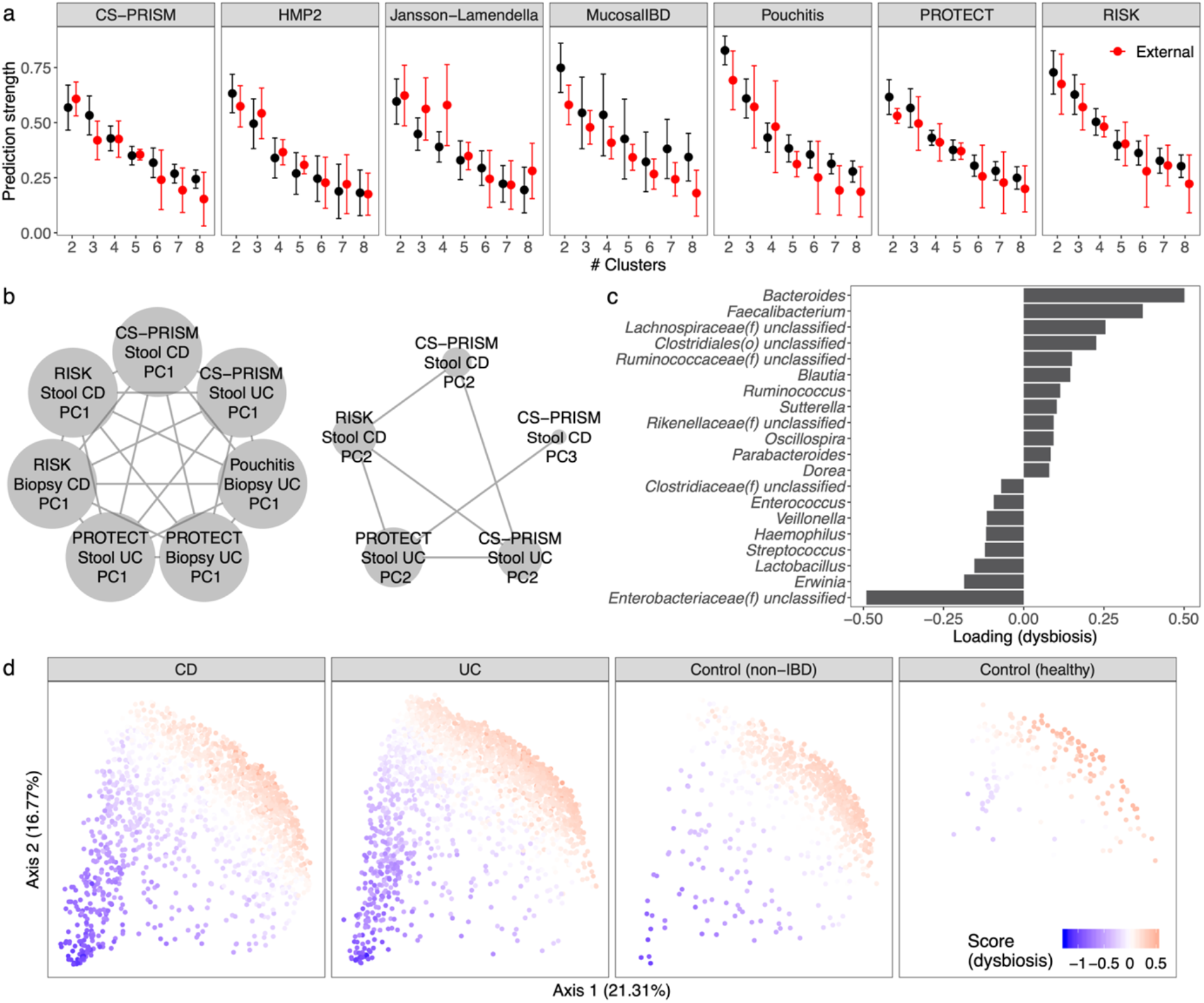
Unsupervised population structure discovery finds no evidence of microbiome-based subtypes in the IBD gut, but a reproducible gradient of continuously variable dysbiosis in disease. **a)** No support was detected for discrete microbiome subtypes (clusters) within the IBD microbiome, neither within cohort nor when evaluated among studies (red bars) using prediction strength^41^. This remained true during stratification within CD and UC, and for additional dissimilarity metric/clustering strength measurements (**Supplemental Fig. 9**). **b)** Conversely, two reproducible, continuously variable patterns of microbiome population structure were identified using groups of similar principal components (**Methods**)^35^. These patterns were consistent within and between cohorts, disease types, and sample types, as well as under different edge strength cutoffs (**Supplemental Fig. 11**), and their consensus loadings were reproducible among cohorts (**Supplemental Fig. 12**). **c)** Top 20 genera with highest absolute loadings for the disease-associated dysbiosis score corresponding to the first cluster in **b**. Many of these taxa were also IBD-associated (**Fig. 3b**). **d**) Distribution of the dysbiosis pattern across CD, UC, non-IBD control, and healthy populations. Although it was defined in an unsupervised way solely within the IBD population, across which the pattern is highly variable, it also differentiates well between IBD and control populations (**Supplemental Fig. 13**).

Conversely, we identified two consistent, continuously varying gradients of microbial community variation in the IBD microbiome (**Fig. 5b-d, Supplemental Fig. 10**). These gradients represent patterns of microbes that occur with greater or lesser abundance in tandem, and which covary across subjects in a population; they were identified as principal component (PC) vectors that recur among different cohorts (see **Methods**)^35^. Briefly, we used the four largest IBD cohorts (CS-PRISM, Pouchitis, PROTECT, and RISK) as training datasets to identify two clusters of consistent PCs (**Fig. 5b**), which were confirmed with sensitivity analysis (**Supplemental Fig. 11**) and validated in the remaining cohorts (**Supplemental Fig. 12**). The consensus loadings (i.e. within-cluster average) representing these two clusters (**Fig. 5c, Supplemental Fig. 10, Supplemental Table 7**) were used to assign continuously varying scores to the IBD population that capture gradient changes in the microbiome that occurred consistently within IBD, across diseases, sample types, and cohorts. This disease-linked “type” of microbiome variation corresponded roughly to severity or extent of inflammation, as detailed below.

In particular, while the second continuous population structure captured the Firmicutes-Bacteroidetes tradeoff present in most gut microbiome studies (**Supplemental Fig. 10**)^9,26,40^, the first continuous score was IBD-specific and corresponded roughly to more extreme disease-associated dysbiosis in CD and UC populations (**Fig. 5d**). This is evidenced by the taxa with highest weights in the scores’ consensus loading vector (**Fig. 5c**), which included taxa differentially abundant between IBD and control populations (**Fig. 3**). The score was consistent both within CD and UC while also further differentiating IBD, non-IBD control, and healthy populations (**Fig. 5d, Supplemental Fig. 13**), even though it was identified unsupervisedly only from diseased subsets. The composition of the score and its population structure are also consistent with our recent definition of dysbiotic gut microbiome configurations corresponding with multi’omic perturbations during IBD activity^9^. Together with the supervised meta-analysis results above, these unsupervised population structure findings confirm that there are no detectable discrete subtypes of the gut microbiome in IBD even among ∼5,000 combined samples, while showing a single continuously variable gradient of microbiome changes reproducibly present during more dysbiotic diseases.

## Discussion

Here, we provide a novel framework for microbial community meta-analysis and apply it to the first large-scale integration of over 5,100 amplicon profiles of the stool and mucosal microbiomes in IBD. This identified a significantly reproducible gradient in the gut microbiome indicative of increasing dysbiosis in subsets of patients. The study also showed no evidence of additional population structure, such as microbiome-driven discrete disease subtypes, within CD or UC. The increased power provided by meta-analysis supported many of the taxonomic associations previously ascribed to IBD (e.g. *Faecalibacterium, Ruminococcus*, Enterobacteriaceae) while uncovering new associations (*Turicibacter, Acinetobacter*) not confidently associated with inflammation by other populations or data types. Almost all effects were exhibited similarly using either stool or mucosal profiling, with a small number of exceptions showing significant differentiation (e.g. *Dehalobacterium*). Novel disease-treatment response interactions were observed (e.g. steroids on Enterobacteriaceae). Finally, the meta-analysis framework developed for the study, MMUPHin, has been extensively evaluated and its performance for batch effect removal, supervised meta-analysis of exposures and covariates, and unsupervised population structure discovery validated on a variety of simulated microbial community types. It is extensible to integration of microbial community taxonomic or functional profiles from other data types (e.g. metagenomic sequencing) or environments.

However, all microbial community meta-analyses should be approached with caution, since in many cases unwanted sources of technical variation between studies (i.e. batch effects) are so large as to potentially mask biological signals even after correction^49-51^ (**Supplemental Notes**). Reducing inter-study variation in microbial community profiles is challenging relative to other ‘omics data types due to 1) the extreme heterogeneity of microbes within most communities (exacerbating both technical and biological differences), and 2) feature zero-inflation arising from both biological and technical reasons^13,52^. Notably, despite these challenges, MMUPHin was able to meta-analyze amplicon profiles in this study both to associate microbial shifts with disease outcome, to associate them with treatment-specific differences, and to identify a single pattern of typical microbial variation within IBD. While previous efforts have developed IBD dysbiosis scores by contrasting patients with control groups^7,9^, this pattern of microbial variation was present specifically within IBD patients (both CD and UC), and in agreement with supervised methods, captured several classes of microbial functional responses in the gut (**Supplemental Note**).

The IBD gut microbiome particularly stands to benefit from meta-analysis, as have other multiply-sampled conditions such as colorectal cancer^53,54^, in order to identify ecological and microbiological changes during the disease that are reproducible across populations. We consider this study based on 16S rRNA gene sequencing to be a proof of concept, able to achieve unprecedented power due to the number of amplicon profiled samples available, but with greater precision possible in future work using e.g. metagenomic and other ‘omics technologies. This also enabled comparison of responses in the stool versus mucosal microbiomes, the latter of which are not amenable to metagenomic profiling from biopsies; these were in overall good agreement, but the few areas of significantly differential responses to inflammation are likely of particular immunological interest. The large sample and population sizes also provide some confidence in ruling out discrete, microbially-driven population subtypes as an explanation for CD and UCs’ clinical heterogeneity. Instead, the work identified a single consistent axis of gradient microbial change corresponding to increasing departures from “normal” microbiome configurations^7,9,55^. This pattern of consistent microbial dysbiosis can continue to be explored in further work on its functional, immunological, and clinical consequences. Overall, this study represents one of the first large-scale, methodologically appropriate, targeted meta-analysis of the IBD microbiome, and the corresponding methodology and its implementation are freely available for future meta-analyses of human-associated and environmental microbial populations.

## Methods

### MMUPHin: a uniform statistical framework for meta-analysis of microbial community studies

We developed MMUPHin (Meta-analysis Methods with a Uniform Pipeline for Heterogeneity in microbiome studies) as a framework for meta-analysis of microbial community studies using taxonomic, functional, or other abundance profiles. It includes components for batch effect adjustment, differential abundance testing, and unsupervised discrete and continuous population structure discovery.

#### Batch adjustment

For microbial community batch correction, we extended the batch correction method developed for gene expression data in ComBat^15^ with an additional component to allow for the zero inflated nature of microbial abundance data. In our model, sample read count*Y* was modelled with respect to both batch variable and biologically relevant covariate(s) *X*:

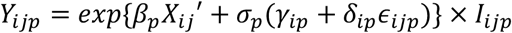

Where *i* indicates batch/study, *j* indicates sample, and *p* indicates feature. *γ*_*ip*_ and *δ*_*ip*_ are batch-specific location and scale parameters. *σ*_*p*_ is a feature-specific standardization factor. *β*_*p*_ are covariate-specific coefficients, and *ϵ*_*ijp*_ is an independent error term following a standard normal distribution. *I*_*ijp*_ is a binary (0, 1) zero-count indicator, to allow for zero inflation of features. As in ComBat, *γ*_*ip*_ and *δ*_*ip*_ are modelled with normal and inverse-gamma priors, respectively. Hyperparameters are estimated with empirical Bayes estimators as in ComBat^15^. The posterior means, 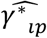 and 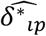, along with standard frequentist estimates 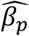 and 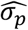 are used to provide batch-corrected count data:

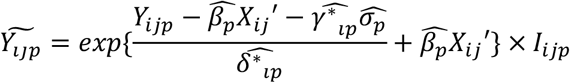

Per-sample feature counts are then re-normalized to keep sample read depth unchanged post-correction. In practice, the user provides sample microbial abundance table (*Y*), batch/study information, and optionally any other covariates *X* that are potentially confounded with batch but encode important biological information. MMUPHin outputs an adjusted profile 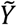 that is corrected for the effect of batches but retains the effects of *X* (if provided).

#### Meta-analysis differential abundance testing

For meta-analytical differential abundance testing, after batch correction, MMUPHin first performs multivariate linear regression within individual studies using previously validated data transformation and modelling combinations appropriate for microbial community profiles (MaAsLin2^33^). This yields study-specific, per-feature differential abundance effects estimations 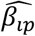, where *i* indicates study and *p* indicates feature. These are then aggregated into meta-analysis effect size with fixed/random effects modelling as implemented in the metafor R package^34^:

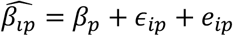

*β*_*p*_ is the overall differential abundance effect of feature *p. ϵ*_*ip*_ is per-study measurement error, and *e*_*ip*_ is study-specific random effects term (not present in fixed-effect models). In practice, the user provides a microbial community profile, study design (batch) information, the main exposure variable of interest, and optional additional covariates. If any meta-analyzed studies include repeated measures (e.g. longitudinal designs), then random covariates can also be provided and will be modelled for such studies. MMUPHin then performs MaAsLin2 regression modelling within each study and aggregates effect sizes of the exposure variable 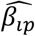 across studies using the resulting random/fixed effects model. The estimated overall effect, 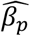, is reported as the overall differential abundance effect for feature *p*.

#### Unsupervised discrete structure discovery

For unsupervised discrete (i.e. cluster) structure discovery of a single study, again after batch correction, MMUPHin uses average prediction strength^41^, an established clustering strength metric, to measure the existence of reproducible clusters among meta-analyzed datasets. Briefly, for each individual dataset, the metric randomly and iteratively divides samples into “training” and “validation” subsets. In each iteration, clustering is first performed on the training samples, across a range of cluster numbers *k*, yielding (for a specific *k*) training sample clusters *A*_*k*1_, *A*_*k*2_, …, *A* _*kk*_. Note that *A*_*k*1_, *A*_*k*2_, …, *A* _*kk*_ jointly forms a partition of the testing sample indices. The same clustering analysis is then performed on the validation samples, and the resulting partition of sample space provides classification membership potentially different from clustering memberships *A*_*k*1_, *A*_*k*2_, …, *A*_*kk*_. Prediction strength for *k*clusters is defined as

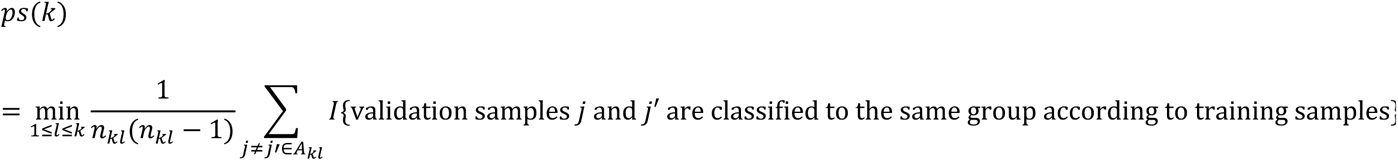

i.e. the minimum (across validation clusters) proportion of same-cluster sample pairs also being classified as the same group by training samples. *n*_*kl*_ = |*A*_*kl*_ |, or the number of test samples in the *l*th cluster.

Average prediction strength is the average of prediction strengths across randomization iterations. Intuitively, it characterizes the degree of agreement between the clustering structures in randomly partitioned validation and training subsets; if *k* is appropriately describing the true number of discrete clusters in the dataset, then average prediction strength should be close to one (training and validation samples agree most of the time).

We additionally generalized this metric to meta-analysis settings, where we aimed to quantify the agreement of clustering structures between studies. In the meta-analytical setting, generalized prediction strength for cluster number *k*in study *i*with validation study *i*′ is

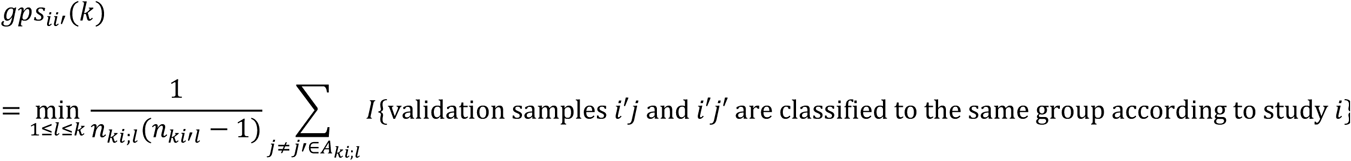

Where *A* _*kil*_ indicates the *l*-th cluster membership in study *i*, when cluster number is specified as *k*; *n* _*kil*_ = |*A* _*kil*_ |. The average generalized prediction in study *i* for cluster number *k* is then defined as the average of *gps*_*ii*′_(*k*) across all *i*′ ≠ *i*, i.e., all validation studies (instead of iterations of randomized partitions). Similar to the single study prediction strength, it describes the generalizability of clustering structure in study *i* in external validation studies.

#### Unsupervised continuous structure discovery

We extended our previous work in cancer gene expression subtyping^35^ to perform unsupervised continuous structure discovery in microbial community profiles. Complementary to discrete cluster discovery, the goal is to identify strong feature covariation signals (gradients) that are reproducible across studies. This is carried out by performing principal component analysis individually in microbiome studies and constructing a network of correlated PCA loading vectors, to identify loadings that are consistently present across studies. In detail, given a collection of training microbial abundance datasets, our method takes the following steps (visualized in **Supplemental Fig. 4**):

1. For each dataset *i*, PCA is performed on normalized and arcsin square root-transformed microbial abundance data. Given a user-specified threshold on variance explained, we record its top PC loading vectors, 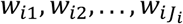, where *J*_*i*_ is the smallest number of top loading vectors that jointly explain percentage of variability in the dataset past a customizable threshold 0 < *threshold*_*v*_ < 1 (default to 80%).
2. For two PC loadings from different datasets *w*_*ij*_ and *w*_*i*′*j*′_, similarity is quantified with the absolute value of cosine coefficient^56^ |*cos* < *w*_*ij*_, *w* _*i*′*j*′_ > |. This yields a network of PC loading vectors associated by weighted edges*w*_*ij*_ and *w* _*i*′*j*′_, retaining edges only if their weight surpasses a customizable similarity threshold (|*cos* < *w*_*ij*_, *w* _*i*′*j*′_ > | > *threshold*_*s*_, 0 < *threshold*_*s*_<1).
3. In the resulting network, we perform community detection^57^ to identify densely connected modules of PCs. Each module by definition consists of PCs from different datasets that are similar to each other - whether or not they occur in the same order or with similar percent variance explained - and which thu represent strong feature covariation signals that are recurrent in studies.
4. For a module *k* containing PC set *M*_*k*_, its consensus vector *W*_*k*_ is calculated as the average of sign-corrected loading vectors in *M*_*k*_, i.e.,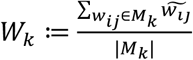. Note that the average is taken not over the original loading vectors *w*_*ij*_, but rather their sign-corrected versions 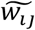. Specifically, the signs of each *w*_*ij*_ in *M*_*k*_ are corrected so that all of the loading vectors have positive cosine coefficients.
5. The module-wide consensus vectors *W*_*k*_ represent strong, mutually independent, and reproducible covariation signals across the microbial datasets; they are used to identify continuously varying gradients in microbial abundance profiles that represent reproducible population structures. Specifically, given a sample with normalized and transformed microbial abundance measurements *x*, its continuous score for module *k* is defined as *x*′*W*_*k*_, as in regular PCA.
6. If additional studies are available, the reproducibility of each *W*_*k*_can be further examined by correlating *W*_*k*_ with the top PC loadings in each such validation study. For each additional study, *W*_*k*_ is considered to be validated in that dataset if its absolute cosine coefficient with at least one of the dataset’s top PCs surpasses the coefficient similarity cutoff *threshold*_*s*_; the number of top PCs to consider in the validation dataset loadings is determined with the same cutoff *threshold*_*v*_.

### Simulation validation of MMUPHin

We performed extensive simulation studies (**Fig. 2, Supplemental Fig. 5-8, Supplemental Table 2**) to validate the performance of each component of MMUPHin. In all cases these employed realistic microbial abundance profiles generated using SparseDOSSA (http://huttenhower.sph.harvard.edu/sparsedossa). This is a model of microbial community structure using a set of zero-inflated log-normal distributions fit to selected training data, in this case drawn from the IBD gut microbiome^6^. Controlled microbial associations with simulated covariates can then (optionally) be spiked in. Note that although the assumed null distributions in MMUPHin and SparseDOSSA are the same (zero-inflated log normal), the models of effects for batch and biological variables are substantially different: MMUPHin assumes exponentiated effects, while SparseDOSSA assumes re-standardized linear effects.

Specifically, SparseDOSSA models null microbial feature abundances using a zero-inflated log-normal distribution:

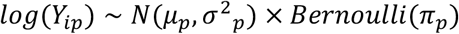

This is the same initial distributional assumption as the MMUPHin batch correction model, when there are no batch or covariates effects. However, for spiked-in associations with metadata (batch, biological variables, etc.), SparseDOSSA uses a different model. Given a simulated, pre-spiking-in feature count vector *Y*_*p*_ with mean *μ*_*p*_^*Y*^ and standard error *σ*_*p*_^*Y*^, as well as a metadata variable vector *X* with mean *μ*^*X*^ and standard error *σ*^*X*^, the post-spiked-in feature count is set to:

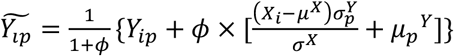

where *ϕ* is a configurable spike-in strength parameter. By this definition, microbial features post-spike-in have the same mean and approximately the same variance as before, the only difference being the added association with the metadata variable(s) used. This is to ensure the counts of the modified feature are not dominated by the values of the target covariate, but instead distributed similarly to real data. The SparseDOSSA association model thus differs from MMUPHin’s model in two substantial ways: i) MMUPHin’s associations are defined within the exponentiated component and are thus better described as a multiplicative effect, whereas SparseDOSSA’s effects are directly applied on untransformed data, and ii) SparseDOSSA additionally ensures realistic data generation with the re-standardization procedure.

Thus, the only component of the SparseDOSSA model that requires fitting to training data is the aforementioned zero-inflated log-normal null distribution. In our analysis, this was always PRISM^6^, while other parameters were specified across a wide range of combinations to simulate different application scenarios. These include the effect sizes of the associated batch and biological variables (i.e. the *ϕ* parameter), number of batches, sample sizes, as well as dimensionality (both the total number of features and the percentage of features randomized to be associated with batch/biological variables). For each combination of simulation parameters, we performed 20 random replications (i.e. running simulation/evaluation with the same parameters but different random seeds). **Supplemental Table 2** presents the full list of parameter combinations.

#### Evaluating batch adjustment

For evaluation of MMUPHin’s batch effect adjustment component, we simulated metadata that included batch (with varying total batch numbers 2, 4, 6, 8), a binary positive control (simulated “biological” covariate), continuous positive control (“biological”), and negative control (binary, and guaranteed to be non-associated with microbial features) variables. Microbial abundance data was simulated to be associated with the batch and the two positive control variables at varying effect sizes (1, 2, 5, 10 for batch variable and fixed at 10 for positive control variables), but not with the negative control variable. We additionally varied the number of samples per batch (20 to simulate multiple-batches in a single study scenario, 100 to simulate meta-analysis with moderate sized studies and 500 to simulate large meta-analysis), total number of microbial features (n=200 and 1000), as well as the percentage of features associated with metadata (5%, 10%, and 20%) (**Supplemental Table 2**).

Performance of batch correction methods was quantified by omnibus associations (PERMANOVA R2) between the simulated microbial abundance data with the batch and positive control variables, before and after batch correction. For ComBat^15^ and our method, batch correction was performed with both positive control variables as well as the negative control variable as covariates. MMUPHin successfully reduced the confounding batch effect, but retained the effect of positive control variables, and did not inflate the effect of negative control variable (**Fig. 2a, Supplemental Fig. 5**).

#### Evaluating meta-analytic differential abundance testing

We evaluated false positive rates (FPR) in particular for meta-analytic feature association testing, specifically the null case in which there are no associations between microbial features and covariates, but false associations can arise in the presence of batch effects with unbalanced distribution of covariate values across studies (**Fig. 2b**). For simulation, we generated a binary covariate unevenly distributed between two “studies” at varying levels of disparity (**Supplemental Table 2**). Microbial abundance data was simulated to be associated only with the two studies and not with the covariate (i.e. study confounded null data), with varying strengths of batch effect (from 0 to 10). The number of samples per batch varied between 100 and 500 to, again, simulate moderate- and large-sized meta-analysis. Lastly, we varied total number of microbial features and the percentage of features associated with metadata as above.

FPRs were calculated as the percentage of simulated microbial features with nominal p-values < 0.05 for associations with the exposure variable. Four data normalization and analysis regimes were evaluated (**Fig. 2c, Supplemental Fig. 6**): a) naive MaAsLin2 model on the study effect confounded null data (without explicitly modelling the batches), b) the quantile normalization procedure, paired with two-tailed Wilcoxon tests, as proposed in ^18^, c) BDMMA as proposed in ^19^, with the default 1,0000 total MCMC sampling and 5,000 burn-in, d) the complete MMUPHin meta-analysis model for the batch corrected data as described above. Note that due to its computational cost we were only able to evaluate the Dirichlet-multinomial regression model on a subset of parameter combinations, namely number of samples per batch = 100, number of features = 200, and percent of associated microbes = 5%. These parameters roughly agree with those used in the simulation analysis in the method’s original publication^19^.

We also evaluated the computational costs of quantile normalization, BDMMA, and MMUPHin (**Supplemental Fig. 6**). For this, the same subset of 20 replications (batch effect 0, exposure imbalance 0, number of samples per batch 100, and number of features 200) were ran through the three methods under the same computation environment (single core Intel(R) Xeon(R) CPU E5-2680 v2 @ 2.80GHz).

#### Evaluating unsupervised discrete structure discovery

To simulate microbial abundance data with known discrete clustering structure, we again used the simulation model above, with microbial feature associations added both with a discrete “batch” variable and a discrete clustering variable, at varying number of batches (2, 4, 6, 8), number of clusters (3, 4, 5, 6), as well as effect size of association (0 to 10 for batch, fixed at 10 for cluster). For the evaluation of MMUPHin’s unsupervised methods (both here and during continuous population structure discovery below), we fixed the number of samples per batch at 500, the number of total features at 1,000, and the percent of associated features at 20%. These were guided by the fact that the underlying unsupervised methods (clustering, PCA) require larger sample sizes for good performance even without batch confounding, and are generally only practical with higher feature dimensions (**Supplemental Table 2**).

Performance of clustering was evaluated as the percentage of replicates in which the right number of synthetically defined underlying clusters was identified using prediction strength, across technical replicates for a fixed combination of simulation parameters. That is, the number of clusters within a simulation was identified as that which maximized prediction strength. This was compared to the “truth” (i.e. the known simulation parameter) and counted as a success only if the two agreed. The percentage of success for a given parameter combination across the 20 random replications was used as the evaluation metric for model performance. We compared the performance of clustering before and after MMUPHin batch correction (**Fig. 2e, Supplemental Table 7**). Note that batch correction is modelled only using the batch variable and specifically not including the cluster variable as a covariate in the batch correction model above, as the underlying cluster structure is unknown in non-synthetic unsupervised analyses settings.

#### Evaluating unsupervised continuous structure discovery

To simulate microbial abundance data with known continuously variable population structure, we spiked in feature associations with both a simulated batch covariate (4, 6, 8) and a continuously varying gradient (uniformly distributed between −1 and 1), at varying number of batches and effect size of both associations (as above). The number of samples per batch, total number of microbial features, and the percentage of features associated were fixed at the same values as above (**Supplemental Table 2**).

Performance of continuous structure discovery analysis was evaluated as the Spearman correlation between the known simulated gradient score and the strongest continuously valued population structure as identified by MMUPHin’s continuous structure discovery method (above). We again compared the performance of continuous score discovery on the batch confounded and batch corrected data (**Fig. 2g, Supplemental Fig. 8**). Note that, as above, batch correction is again modelled only using the batch variable and does not have any access to the synthetic continuous gradient, as any underlying continuous population structure is unknown during unsupervised analyses settings.

### Collection and uniform processing of ten IBD microbiome studies employing 16S rRNA gene sequencing

#### Study inclusion and raw sequence data

We curated 10 published 16S rRNA gene sequencing (abbreviated 16S) gut microbiome studies of IBD for meta-analysis (**Table 1, Supplemental Table 1**). Demultiplexed raw sequences were either downloaded from EBI (Jansson-Lamendella and Herfarth) or available locally as previously generated (other eight studies). Metadata were obtained either directly from the sequence repository/manuscript (Herfarth, Jasson-Lamendella, HMP2, MucosalIBD, PROTECT, RISK), or from collaborators (BIDMC-FMT, CS-PRISM, LSS-PRISM, Pouchitis). This resulted in a total of 5,151 samples and 2,179 subjects available prior to processing and quality control.

#### Metadata curation

We manually curated subject- and sample-specific metadata across studies to ensure consistency. Variables collected and curated include:

- Disease (CD, UC, control), universally available.
- Type of controls (non-IBD, healthy). Control information was available directly for CS-PRISM, Jansson-Lamendella, and Pouchitis, inferred from study design described in manuscript for Herfarth, HMP2, MucosalIBD, and RISK (all non-IBD controls), and not applicable for BIDMC-FMT, LSS-PRISM, and PROTECT (only has IBD subjects).
- Sample type (biopsy, stool), universally available.
- Body site of biopsy sample collection (ileum, colon, rectum), with more detailed classifications recorded separately in case of need. Mappings for the relevant datasets are:
  ∘ CS-PRISM: terminal ileum, neo-ileum, pouch are aggregated as ileum; cecum, ascending/left-sided colon, transverse colon, descending/right-sided colon, sigmoid colon were aggregated as colon; rectum classification was kept unchanged.
  ∘ HMP2: ileum classification kept unchanged; cecum, ascending/right-sided colon, transverse colon, descending/left-sided colon, and sigmoid colon were aggregated as colon.
  ∘ MucosalIBD: all terminal ileum samples, aggregated to ileum.
  ∘ Pouchitis: terminal ileum, pouch, pre-pouch ileum aggregated as ileum; sigmoid colon aggregated to colon.
  ∘ PROTECT: all rectum samples, classification kept unchanged.
  ∘ RISK: terminal ileum was aggregated to ileum; rectum kept unchanged.
- Montreal classifications:
  ∘ Location for CDs (L1, L2, L3, and possible combinations), available for BIDMC-FMT, CS-PRISM, Herfarth, Jansson-Lamendella, LSS-PRISM, and Pouchitis.
  ∘ Behavior for CDs (B1, B2, and B3), available for CS-PRISM, Herfath, Jansson-Lamendella, LSS-PRISM, Pouchitis, and RISK.
  ∘ Extent for UCs (E1, E2, and E3), available for CS-PRISM, Jansson-Lamendella, LSS-PRISM, Pouchitis, and PROTECT.
- Age at sample collection (in years), available for BIDMC-FMT, CS-PRISM, Herfarth, HMP2, LSS-PRISM, MucosalIBD, Pouchitis, PROTECT, RISK.
- Age at diagnosis (in years). Directly available for CS-PRISM, HMP2, LSS-PRISM, and Pouchitis, inferred as baseline age for PROTECT and RISK as these were new-onset cohorts.
- Race (White, African American, Asian / Pacific Islander, Native American, more than one race, others). Directly available for CS-PRISM, Herfarth, HMP2, PROTECT, and RISK, inferred from manuscript cohort description for Jansson-Lamendella (all Caucasian cohort).
- Gender (male/female). Available for BIDMC-FMT, CS-PRISM, Herfarth, HMP2, Jansson-Lamendella, LSS-PRISM, MucosalIBD, Pouchitis, PROTECT,
- Treatment variables, including antibiotics, immunosuppressants, steroids, and 5-ASA. These variables were encoded as yes/no to indicate, approximately, currently receiving them at the time of sampling. Additional information such as specific medication or delivery method was recorded separately if available in case of need. We note the potentially confounding difference in studies’ definitions of treatment: for Pouchitis and PROTECT authors defined antibiotics as receiving the treatment within the past month (30 days for Pouchitis, 27 days for PROTECT), whereas for CS-PRISM, HMP2, LSS-PRISM, and RISK such determination was not possible (antibiotics “yes” was defined as “currently taking”). Likewise, we had no additional information to determine the time extent for the other three treatments, beyond that according to metadata/publication, patients were “currently taking” the treatment at sample collection.

For a comprehensive list of curation mapping schema, please refer to our metadata curation repository: https://github.com/biobakery/ibd_meta_analysis.

#### 16S amplicon sequence bioinformatics and taxonomic profiling

Sequences were processed, per-cohort, with the published, standardized bioBakery workflow^58^ using the UPARSE protocol^59^ (version v9.0.2132-64bit). For all studies, demultiplexed sequences were truncated at 200bp max length and filtered by maximum expected error of one^59^. Operational taxonomic units (OTUs) were clustered at 97% identity and aligned using USEARCH with 97% identity to the Greengenes database 97% reference OTUs (version 13.8)^60^ for taxonomy assignment. The resulting Greengenes identifiers for OTUs were used as basis for matching features (taxa) among cohorts.

#### Quality control

Across samples, a median of 81.51% reads / sample passed quality control filtering and were successfully assigned to OTUs with Greengenes identifiers (**Supplemental Fig. 1**). These 8,921 raw OTUs aggregated to a total of 1,122 genera prior to quality control. We retained taxa that exceeded 5e-5 relative abundance with at least 10% prevalent in at least one study; this criterion generally removes spurious OTU assignments while retaining rare organisms if confidently present in at least one study. Lastly, we also removed low read depth samples with less than 3,000 total sequences, which retained 78.34%-100% samples per cohorts (**Supplemental Table 1**). The final resulting taxonomic profile, used for all further analysis, aggregated into 249 total genera spanning 4,789 samples (OTUs unclassified under a particular taxonomy level were aggregated as “unclassified” feature under that taxon, e.g. “Enterbacteriaceae unclassified” accumulates all OTUs’ abundances under the family that could not be classified at the genus level.

#### Data availability

Quality controlled (truncated and filtered) sequences, Greengenes mapped OTU count profiles, and curated sample metadata are available at the Human Microbial Bioactives Resource Portal (http://portal.microbiome-bioactives.org).

### Applying MMUPHin to IBD gut microbiome meta-analysis

For the resulting collection of microbiome studies, batch and study effects was performed using MMUPHin on both the genus level feature abundance profiles. For either taxonomic rank, batch (i.e., sequencing run) effect correction was first performed within individual studies (when batch/plate information was available, applicable to BIDMC-FMT, CS-PRISM, LSS-PRISM, MucosalIBD, and RISK). Microbial abundance profiles across all studies were then jointly corrected for study effects, while modelling disease status (IBD or control), disease (CD or UC), and sample type (biopsy or stool) as covariates. Reduction of batch and study effects was evaluated by PERMANOVA R2 (**Fig. 3a**).

### Association analyses

#### Omnibus testing of microbial composition associations

We used PERMANOVA tests (2,000 permutations) as implemented in the R package vegan^37^ using Bray-Curtis dissimilarities for all omnibus association tests of overall microbial community structure with covariates (**Fig. 3a**). Where appropriate, R2s were calculated conditioning on the necessary covariates; specifically, CD/UC Montreal classifications were conditional on CD/UC samples respectively, treatment was conditional on IBD status, biopsy location was conditional on a sample being a biopsy, and all covariates were conditional on being non-missing. Otherwise, variables were tested marginally (that is, each as the sole variable in the model). Importantly, to account for repeated measures within subjects for longitudinal studies, we adopted the blocked permutation strategy as in ^9^, where per-sample measurements (sample type, biopsy location, treatment) were permuted within subjects, and per-subject measurements (disease, demographics) were permuted along with subjects (but within cohorts, relevant for the all-cohorts evaluation). For a full list of the model and permutation strategies that this resulted in for our analysis, please refer to **Supplemental Table 3**. Finally, per-variable p-values were adjusted with Benjamini-Hochberg false discovery rate control on a per-study basis.

#### Per-feature meta-analysis differential abundance testing

To identify microbial features individually significantly associated with one or more covariates, we applied MMUPHin’s differential abundance testing model as described above. Cohorts were first stratified by sample type (biopsy or stool) and, where appropriate, diseases (CD or UC) prior to model fitting. Arcsin square root-transformed genus level taxon abundances were tested for covariate associations in individual cohort strata with multivariate linear modelling (linear random intercept model adopted for longitudinal studies). Covariates used for adjustment include age, gender, and race for disease variables, and additionally disease status for treatment variables. Effect sizes across cohort strata were aggregated with a random effects model with restricted maximum likelihood estimation^34^. P-values were FDR adjusted across features for each variable. For the full list of models adopted as well as cohort stratification strategy, please refer to **Supplemental Table 3. Fig. 3b** visualizes the aggregated meta-analysis effects; for individual study results refer to **Supplemental Table 4**.

#### Testing for phenotypic severity within CD and UC patients

Meta-analytical testing of features associated with CD behavior and UC extent classifications were performed with similar models (**Supplemental Table 3**). Specifically, within each study’s CD patients, the tests for contrasts B2 versus B1 and B3 versus B1 are performed by

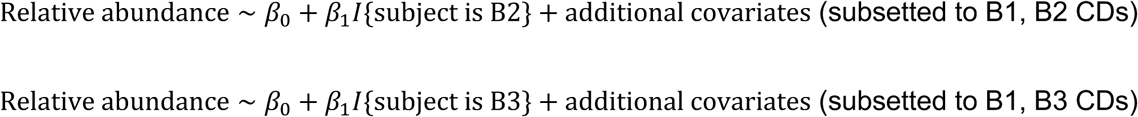

The two *β*_1_ coefficients, once aggregated with meta-analysis, were reported as the effect sizes shown in **Fig. 4a**, along with their FDR corrected q-values (adjusted across features for each test).

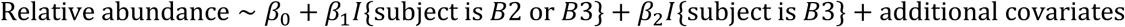

*β*_2_ in this model corresponds to the effect of B3 in addition to the overall contrasts between B23 versus B1. The meta-analysis aggregated p-values of these effects were reported as the differentiation between the most severe and “medium” severity phenotypes (vertical bars indicating significance in **Fig. 4a**). Note that FDR adjustment of this effect was performed across the subset of features with at least either B2 versus B1 or B3 versus B1 effect significant (i.e., the subset of features visualized in **Fig. 4a**). Equivalent models were adopted for contrasts between extent categories of UC patients. Individual study results for the aggregated effects in **Fig. 4a** are in **Supplemental Table 5**.

#### Interaction effects testing

To test for interaction effects with sample type and diseases, we fit meta-analysis moderator models^34^ on the per cohort strata effects:

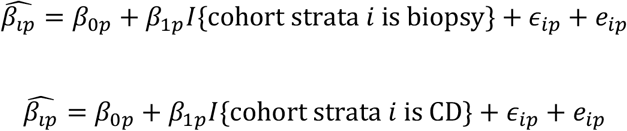

The moderator effects *β* _1*p*_ correspond to the interaction effect between the exposure under evaluation (disease, treatment, etc.) with the moderator variable. **Fig. 4b** visualizes the two example features, *Dehalobacterium* and Enterobacteriacea; al significant interactions as well as individual study effects are in **Supplemental Table 6**.

### Population structure analyses

#### Discrete structure discovery

We performed discrete subtype discovery (i.e. “enterotyping”^61^) in IBD, CD, and UC populations across studies (longitudinal studies subsetted to baseline samples), using MMUPHin’s discrete structure discovery component. Only studies with at least 33 samples were considered for clustering analysis, as this was the sample size in the original enterotype paper^26^. Specifically, clustering was performed on Bray-Curtis dissimilarity by the partition-around-medoid method as implemented in R package cluster; the same method was adopted in previous enterotyping efforts including the original enterotype paper^26,40^. Clustering was evaluated with prediction strength and validated externally with MMUPHin’s generalized prediction strength as described above. Across studies, we found no evidence to support a particular number of clusters within IBD, CD, or UC populations (**Fig. 5a, Supplemental Fig. 9**), suggesting that the IBD microbiome does not have discrete clusters.

We additionally extended our clustering evaluation analysis to other dissimilarity metrics (Jaccard, root Jensen-Shannon divergence) and clustering strength measurements (Calinski-Harabasz index, average silhouette width), which were also explored in previous efforts^40^, Importantly, the original enterotype paper adopted root Jensen-Shannon divergence and Calinski-Harabasz index for cluster discovery. Across combinations of these additional dissimilarities and clustering strength metrics, we also found no evidence to support discrete clusters (**Supplemental Fig. 9**).

#### Continuous structure discovery

Continuous structure discovery was performed with MMUPHin’s corresponding component. The four largest studies (CS-PRISM, Pouchitis, PROTECT, RISK) were subsetted to baseline samples (only relevant for PROTECT), stratified by CD/UC and biopsy/stool sample type, and used as the training sets for MMUPHin. The minimum variance explained threshold (*threshold*_*v*_) was set to default (80%), but we varied the PC similarity (evaluated by absolute cosine coefficient) cutoff*threshold*_*s*_ between 0.5 and 0.8 to assess the sensitivity of the two identified PC clusters in **Fig. 5b** (corresponding to *threshold*_*s*_ = 0.65). As we show in **Supplemental Fig. 11**, with a small *threshold*_*s*_(0.5) PC networks become denser, with the two PC clusters in **Fig. 4b** forming key components of two larger clusters; when *threshold*_*s*_ is large (0.8) the network is sparser, with only the most highly similar nodes of the two clusters forming smaller communities. We thus concluded that the two identified clusters in **Fig. 5b** were not sensitive to the cosine coefficient threshold, as they were recurrently identified in both smaller and larger cutoff scenarios.

#### Continuous structure validation

We validated the consistency of the two clusters’ corresponding continuous scores in all IBD cohorts, non-IBD and healthy control samples, as well as a randomly permuted mock study (as a negative control). The reproducibility of each continuous score within a study was defined as the maximum absolute cosine coefficient between the score’s consensus loading (as provided by MMUPHin) and the top three principal component loadings discovered independently within that study. Note that the number of top principal components considered here was set to a fixed value (three) instead of based on a percent variance cutoff as in the MMUPHin continuous structure discovery stage. This is because in the two identified clusters in **Fig. 5c**, the latest included node was PC3. The randomly permuted study consisted of 473 samples (median validation data sets sample size) randomly selected from the entire meta-analysis collection, but each sample’s microbial abundance was independently permuted across features. This was to simulate a “negative control” dataset where there should be no continuous population structures.

As we show in **Supplemental Fig. 12**, the dysbiosis score was well validated across studies, except for healthy control samples and the negative control dataset. The Firmicutes-versus-Bacteroidetes trade-off score, on the other hand, was reasonably well reproduced in all studies and particularly well-established in healthy samples, but, again, was not significantly detected in the negative control dataset.

#### Continuous score assignment

Assignment of continuous scores was straightforward given the two consensus loading vectors provided by MMUPHin. Within each study, arcsin square root-transformed relative abundances were centered per-feature, the transformed abundance matrix was then multiplied by each consensus loading via dot product to generate per-sample continuous scores. These scores were used for visualization as in **Fig. 4d** and **Supplemental Fig. 10**, as well as for testing the difference between CD, UC, non-IBD, and healthy control populations as in **Supplemental Fig. 13** We provide the two consensus loadings in **Supplemental Table 7;** interested researchers can follow these steps to assign the two continuous scores in other datasets.

