## Supplementary Figures, Tables, and Notes for "Population Structure Discovery in Meta-Analyzed Microbial Communities and Inflammatory Bowel Disease": Supp_Material_Captions.docx

### Supplemental Material Captions

**Supplemental Fig. 1: Read depth varies across studies and is correlated with number of detected taxonomic feature. a)** Per-sample total read depth varies across studies, and vast majority of reads are successfully classified with our bioinformatics protocols. **b)** Median read depth if each study is correlated with number of detected microbial features, both before and after prevalence and abundance filtering.

**Supplemental Fig. 2: Globally prevalent taxa vary in study-specific mean abundance and prevalence, jointly affected by biological and technical factors**. **a)** For genera that are highly prevalent (>70% overall prevalence), per-study mean abundance can vary greatly. Each point represents a study-specific mean abundance of one feature. **b)** For other globally present genera, study-specific prevalence varied (left panel) and often correlated with median read depth (right panel).

**Supplemental Fig. 3: Per-study prevalence of taxa that are not globally present, i.e., missing from only one or a few studies.** Features are grouped by number of present studies.

**Supplemental Fig. 4: MMUPHin constructs correlated principal component loadings for robust continuous structure discovery. a)** Principal component analysis is performed in individual studies, identifying the strongest signals present in each, before **b)** principal component loadings from different studies are compared to each other, identifying loadings that are highly similar to each other. **c)** Clusters of similar PC loadings are identified through network community detection, representing strong, recurrent, continuous variation patterns across multiple studies, which can then be used to assign robust, continuously varying gradients in individual cohorts (**d**).

**Supplemental Fig. 5: Full set of performance evaluation and comparison of MMUPHin's batch adjustment method. a-d** Panels are organized by variables (batch, binary positive control, continuous positive control, and negative control) evaluated by the PERMANOVA R2.

**Supplemental Fig. 6: Full set of performance evaluation and comparison of MMUPHin's meta-analysis differential abundance testing method.** MMUPHin consistently controls false positive rates across different confounding cohort exposure distribution imbalance set up, when compared to naïve regression, quantile normalization, and BDMMA methods (**a-e**). Note that due to computational cost, BDMMA (purple) was only evaluated for the subset of simulation cases most similar to those evaluated in its publication^19^.

**Supplemental Fig. 7: Full set of performance evaluation and comparison of MMUPHin's discrete structure discovery method. a-d** Panels are organized by simulated true number of clusters (3-6).

**Supplemental Fig. 8: Full set of performance evaluation and comparison of MMUPHin's continuous structure discovery method.**

**Supplemental Fig. 9: Comprehensive evaluation provided no evidence to support discrete enterotypes in IBD cohorts.** Combinations of clustering strength evaluation metrics (prediction strength, Calinski-Harabasz index, and average silhouette width), paired with different dissimilarity measures (**a** Bray-Curtis, **b** Jaccard, and **c** square root Jensen-Shannon divergence) were evaluated, with no consistent support for the existence of enterotypes (i.e., “peaking” of clustering strength metric at a particular cluster number $k$) across cohorts.

**Supplemental Fig. 10: Second continuous score identified by MMUPHin characterizes dominant phyla trade-off in the IBD gut. a)** Consensus loading vector corresponding to the second continuous score is dominated, in opposing directions, by genera from the Firmicutes and Bacteroidetes phyla. **b)** The phyla trade-off score is consistently present across diseased and control populations.

**Supplemental Fig. 11: Continuous PC loading clusters are not sensitive to cosine coefficient cutoff.** The same core members of each of the two PC loading clusters are repeatedly observed across different cutoffs (**a-c**).

**Supplemental Fig. 12: The two continuous PC scores are validated in testing cohorts.** The dysbiosis score was reproduced in all training and validation cohorts except for in healthy populations and the negative control dataset (**a**), while the common gut phyla trade-off score was reproduced across most of the diseased and healthy populations except for negative control (**b**).

**Supplemental Fig. 13: The dysbiosis score differentiates between CD, UC, non-IBD control, and healthy populations.** P-value was obtained via two-sample Wilcoxon rank sum tests and adjusted by Benjamini-Hochberg procedure.

**Supplemental Table 1: Additional demographic, clinical, and bioinformatics characteristics of included studies.**

**Supplemental Table 2: Full set of varying simulation parameters evaluated for MMUPHin validation.**

**Supplemental Table 3: Detailed specification for omnibus and per-feature differential abundance testing models.**

**Supplemental Table 4: Per-study and meta-analytically aggregated effect sizes of per-feature differential abundance tests performed.**

**Supplemental Table 5: Per-study and meta-analytically aggregated effect sizes of tests on per-feature abundance associated with disease phenotypic severity.**

**Supplemental Table 6: Per-study and meta-analytically aggregated effect sizes of tests on the interaction between disease/treatment and sample type/disease on per-feature abundances.**

**Supplemental Table 7: Consensus loadings for the two identified continuous scores characterizing the IBD gut microbiome population structure.**
