## Supplementary Figures, Tables, and Notes for "Population Structure Discovery in Meta-Analyzed Microbial Communities and Inflammatory Bowel Disease": Supplemental_Notes.docx

**Per-feature study heterogeneity before MMUPHin processing**

Even upon initial inspection and after uniform processing, a variety of inter-study differences existed to be corrected by MMUPHin. While the majority of genera were present in at least two studies (211 of 249), the genus richness of each study was significantly correlated with its median read depth (Spearman 0.55, one-sided p = 0.049, **Supplemental Fig. 1**). Likewise, among the most prevalent taxa (>70% prevalence overall, present in all studies), study-specific average abundances varied significantly, likely due to a combination of biological and technical factors (**Supplemental Fig. 2**). The Pouchitis study^1^, for example, featured only subjects with j-pouch surgeries, and was uniquely enriched for *Ruminococcus*. PROTECT^2^, in contrast, harbored patients with severe disease and those in remission, allowing us to observe a health-associated enrichment of *Bifidobacterium*. Among the remaining globally prevalent taxa, study-specific prevalence also varied, e.g. in the typically oral taxa *Fusobacterium*, *Porphyromonas*, and *Gemellaceae*^3^, as well as understudied but biologically important clades such as *Christensenellaceae* and *Turicibacter*^4,5^. This heterogeneity was again often associated with read depth (**Supplemental Fig. 2**). Microbes missing from only one or a few studies included common amplicon sequencing contaminants (e.g. *Sphingomonas*, *Burkholderia*)^6^; these were often present at low levels in many but not all protocols (**Supplemental Fig. 3**). Notably, the Jansson-Lamendella study uniquely detected many microbes, likely due to an overall much greater sequencing depth (**Supplemental Fig. 1**). These real-world meta-analysis results echo previous findings on controlled sample sets^7,8^ in which amplicon profiling results in particular could differ strikingly among protocols and populations, despite uniform data handling, suggesting, again, the importance of technical batch effect adjustment prior to downstream analysis.

**Validation of MMUPHin’s supervised differential abundance testing**

As is often noted in GWAS meta-analyses, technical artifacts can cause false associations when confounded with variables of interest^9^; MMUPHin successfully corrected this behavior in simulated microbiome profiles (**Fig. 2c-d**, **Supplemental Fig. 6**). Specifically, we simulated data in which an “exposure” variable was not associated with the microbiome, but was distributed with imbalance between two “batches” (e.g. more cases in one batch, more controls in another). In the presence of nontrivial batch effects, microbial features were spuriously associated with the exposure when appropriate correction and meta-analysis procedures were not applied (**Fig. 2d**). We evaluated the performance of quantile normalization^10^, BDMMA^11^, and MMUPHin on these batch-confounded data, summarized as the proportion of features incorrectly associated with the exposure (false positive rate or FPR, **Methods**). MMUPHin successfully controlled FPR, even in the presence of strong batch effect and exposure distribution imbalance (**Fig. 2c**, **Supplemental Fig. 6**). Quantile normalization conservatively controlled false positives under mild exposure imbalance, but inflated FPRs for greater imbalances; this was because the procedure normalizes case distributions against controls in all batches, and with sufficient imbalance, case-enriched batches have too few control samples to act as a reliable reference distribution. BDMMA has overly conservative FPRs, since the lack of adjustment for zero-inflations generates few positive testing results when microbial features are indeed sparse^11^; it also had greatly increased computational costs relative to other methods (**Supplemental Fig. 6**). Additionally, again, quantile normalization and BDMMA are both also only appropriate for case-control differential abundance tests with at most one covariate. In contrast to existing methods, MMUPHin efficiently enables these as well as any other downstream analysis of microbial community profiles, while consistently controlling false positives in batch-confounded data.

**Confirmation of previously reported IBD-microbial associations through meta-analysis**

Our meta-analysis identified taxa associated with IBD reported in previous findings (**Fig. 3b**), included microbes within the Clostridiales order. Genera from the Lachnospiraceae, Ruminococcaceae, and Christensenellaceae families were depleted in IBD, while the family Veillonellaceae had members both enriched (*Veillonella*, *Dialister*) and depleted (*Phascolarctobacterium*) in disease. Several taxa within the order Bacteroidales were negatively associated with disease, including *Parabacteroides*, *Prevotella*, *Butyricimonas*, and *Alistipes*. Other taxa commonly reported to be enriched for IBD were also observed, including *Fusobacterium*, *Enterococcus*, and *Erwinia*. Finally, several taxa placed with limited resolution were associated with IBD, including unclassified members of the Coriobacteriaceae, Peptococcaceae (both depleted)*,* and Enterobacteriaceae (enriched). Note that the last likely represents *Escherichia coli*, commonly found to be enriched in IBD, but poorly taxonomically resolved by the combination of 16S rRNA gene protocols and taxonomic placement data used here^12^.

**Differential response of the IBD microbiome towards treatment groups**

The similarities between IBD-associated and antibiotic-responsive taxa suggest both residual confounding between disease severity and receipt of antibiotics - although these two variables were analyzed separately - as well as similar microbial physiological sensitivities to immune-driven versus chemical selective pressures. Immunosuppressants, arguably the most extreme type of treatment among these groups, induced microbial responses that were similar in direction to antibiotics, but never statistically significant. This likely speaks to their greater direct effect on host bioactivity without direct microbial community impact (unlike antibiotics). 5-ASAs, mainly administered to UC patients, had opposing effects when compared to antibiotics, immunosuppressants, and steroids, corresponding to enrichments in “anti-inflammatory” taxa. This may speak either to its use in less extreme cases of inflammation, or to a greater success in restoring a less dysbiotic microbial community (as opposed to targeting microbes or inflammatory activity directly).

### MMUPHin’s unsupervised population structure discovery in microbial communities

For discrete structures, MMUPHin utilizes established clustering strength evaluation metric^13^, to a) evaluate the existence of discrete clusters within individual microbiome studies, and b) validate the reproducibility of such structures among studies meta-analytically. For continuous structures, our method generalizes single study principal component analysis (PCA^14^) to multiple studies by constructing a network of correlated top PC loadings^15^, thus identifying major axes of variation that explain the largest amount of heterogeneity between microbial profiles and are also consistent across studies. These unsupervised population structure discovery methods thus complement their supervised counterparts, enabling comprehensive characterization of consistent microbial perturbations after correction for technical artifacts in meta-analyzed cohorts.

MMUPHin’s batch correction successfully improved discrete cluster structure discovery, especially when the numbers of true clusters and confounding batches were both relatively small, but the effect of batch signals was large enough to interfere with clustering structure if uncorrected (**Fig. 2e-f**, **Supplemental Fig. 7**). Clustering performance was evaluated by the success rate of correctly identifying the true number of underlying simulated clusters across randomization replicates (i.e. all simulation parameters held constant, different randomized iterations) using prediction strength^13^ (**Methods**). When uncorrected, batch effects directly impacted the ability of cluster discovery to identify the correct population structure, or even suggested the wrong number of clusters. MMUPHin ameliorated this behavior via batch correction (**Fig. 2e-f**), even though correction in unsupervised analysis cannot directly model a biological signal as it is unknown during the discovery stage. Similarly, continuously variable population structure gradients were better identified in batch-confounded microbial data after MMUPHin correction (**Fig. 2g-h**, **Supplemental Fig. 8**). Continuous structure in microbial data is defined here as traits strongly associated with abundance profiles that are continuous rather than discrete (**Fig. 2h**), such as dominant phyla^16^ or disease-associated dysbiosis^17,18^. Given a collection of microbiome studies, our method identifies top continuous “scores” that consistently characterize the strongest signals of variation across studies using correlated principal component vectors (**Methods**, **Supplemental Fig. 4**). We evaluated the performance of our method by assessing the Spearman correlation between a simulated true underlying continuous structure and the predicted top score (**Methods**). Even when batch strongly confounded the simulated biological signals, MMUPHin was again able to correctly recover the expected continuous structure after batch-normalizing the data.

**Interaction effect of treatment with diseases through meta-analysis**

Specifically, we observed preferential effects of some treatments on individual taxa only within certain diseases (**Supplemental Table 6**). An example of this is the effect of steroids on the Enterobacteriaceae, which tended to be more abundant in CD patients receiving steroids, but less abundant in UC recipients. This could be an effect of route of delivery, since UC patients are more likely to receive essentially topical (enema/foam) steroids, which would have a proportionally greater effect on mucosally-associated microbes such as the Enterobacteriaceae. Alternatively, the more extreme dysbiosis typical of CD may be ecologically resistant to a relatively modest immunomodulatory therapy such as steroids, which will have greatest effect at the mucosa regardless of delivery. Other differences here (**Supplemental Table 6**) may represent secondary effects of the gut microbiota’s biogeographical stratification^19^, e.g. if both disease-local inflammation and different treatment regimes happen to affect microbes that are mucosally or lumenally enriched. More speculatively, treatment-subtype interactions such as steroids and the Enterobacteriaceae may speak to strain-level differences in colonization and subsequent treatment response between CD and UC, which can be better explored when similarly-large shotgun metagenomic meta-analyses are feasible.

**Supplemental discussions**

In our IBD application, the effect of batch and cohort differences was reduced by normalization, but still remained the strongest source of variation across microbial profiles, surpassing even the most prominent biological signals such as inflammation, IBD subtype, or mucosal versus stool differences (**Fig. 3a**). Reducing inter-study variation in microbial community profiles is challenging, relative to other ‘omics data types, for at least two reasons. First, the extreme heterogeneity of microbes within a community, between different community types, and among data generation protocols leads to experimental variation greater than that in areas such as genotyping or RNA-seq^7,20^. Second, statistical methods for correcting this resulting variation are also more difficult to apply accurately, mainly due to the abundance of zero count feature measurements arising both for technical and for biological reasons^21^. Ideally, these two types of zero measurements should be corrected for differently if features are differentially absent across batches/cohorts. However, they are difficult to distinguish post hoc, and it is further non-trivial to re-impute nonzero values for biological zeros (as opposed to adjusting systematic bias in non-zero measurements)^22^. Our meta-analysis framework took a conservative approach in addressing this issue, only correcting the non-zero component of feature counts; features that have low prevalence can borrow power from more prevalent ones through the empirical Bayes modelling framework as provided by ComBat^23^ (**Methods**). Future methodology development would ideally extend this framework both to distinguish between technical and biological zeros, and to - cautiously - impute nonzero values when appropriate.

Notably, while previous efforts have developed IBD dysbiosis scores by contrasting patients with control groups^17,18^, it is intriguing that this study identified essentially the same microbiome variation in an unsupervised way solely by assessing the most consistent patterns of variation within CD and UC populations. In agreement with supervised methods, this dysbiosis axis captures three classes of microbial functional responses: the gain of pro-inflammatory, aerotolerant opportunists (Enterobacteriaceae, *Erwinia*, *Enterococcus*, etc.), many typical of the oral cavity (e.g. *Fusobacterium*, *Veillonella*), at the expense of the gut’s typical SCFA-producing fastidious anaerobes (e.g. Faecalibacterium, Lachnospiraceae, Ruminococcaceae). Importantly, this single, consistent set of changes in the microbiome characterized all IBD populations in our meta-analysis, spanning cross-sectional and longitudinal cohorts as well as CD and UC disease severities and extents. Microbes newly associated with this axis, such as *Turicibacter* and *Acinetobacter*, may share similar physiologies, but might also represent new molecular activities in the disease when functionally characterized in the future.
