## Supplementary figures and images for "Population Structure Discovery in Meta-Analyzed Microbial Communities and Inflammatory Bowel Disease"

### suppFig1.pdf

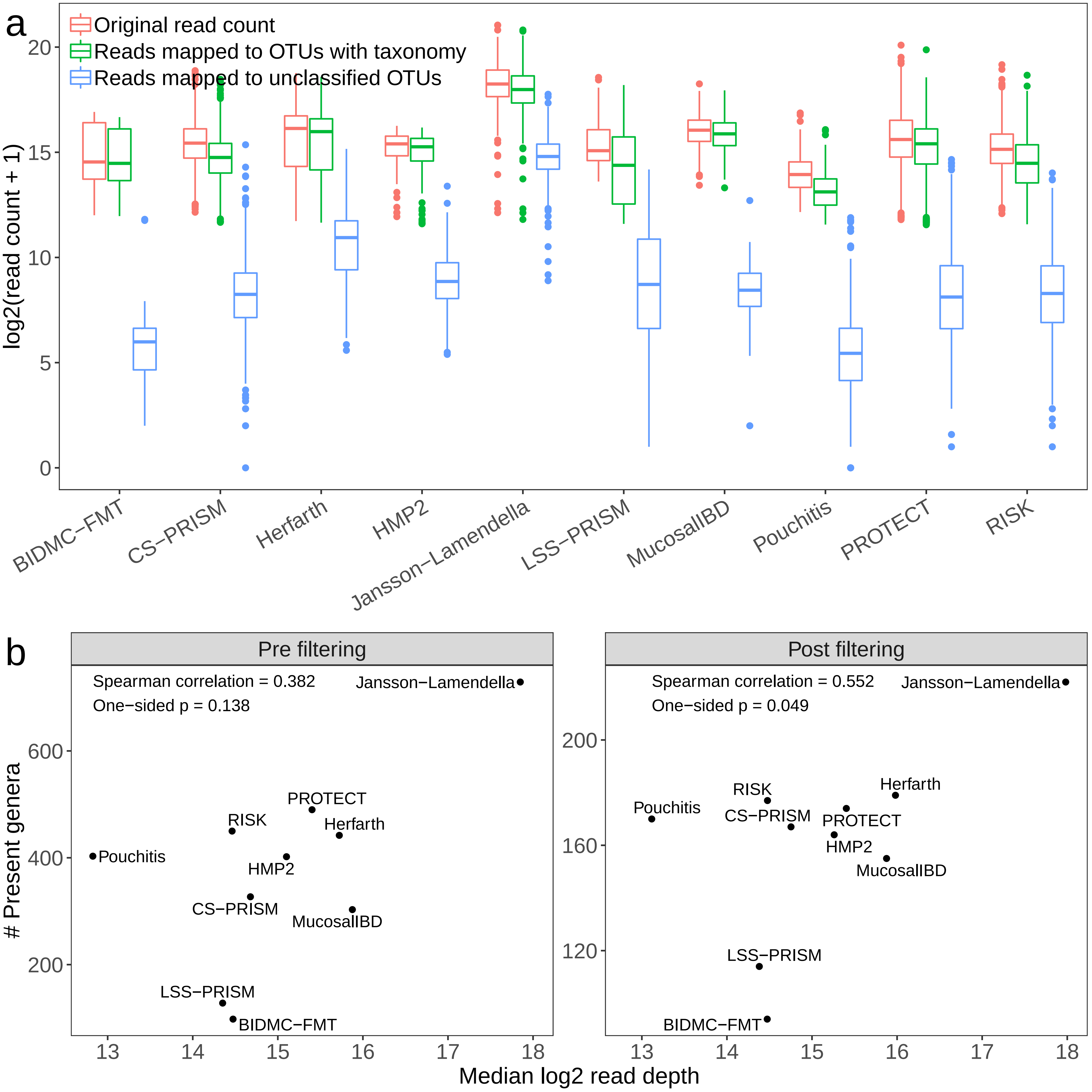

### suppFig2.pdf

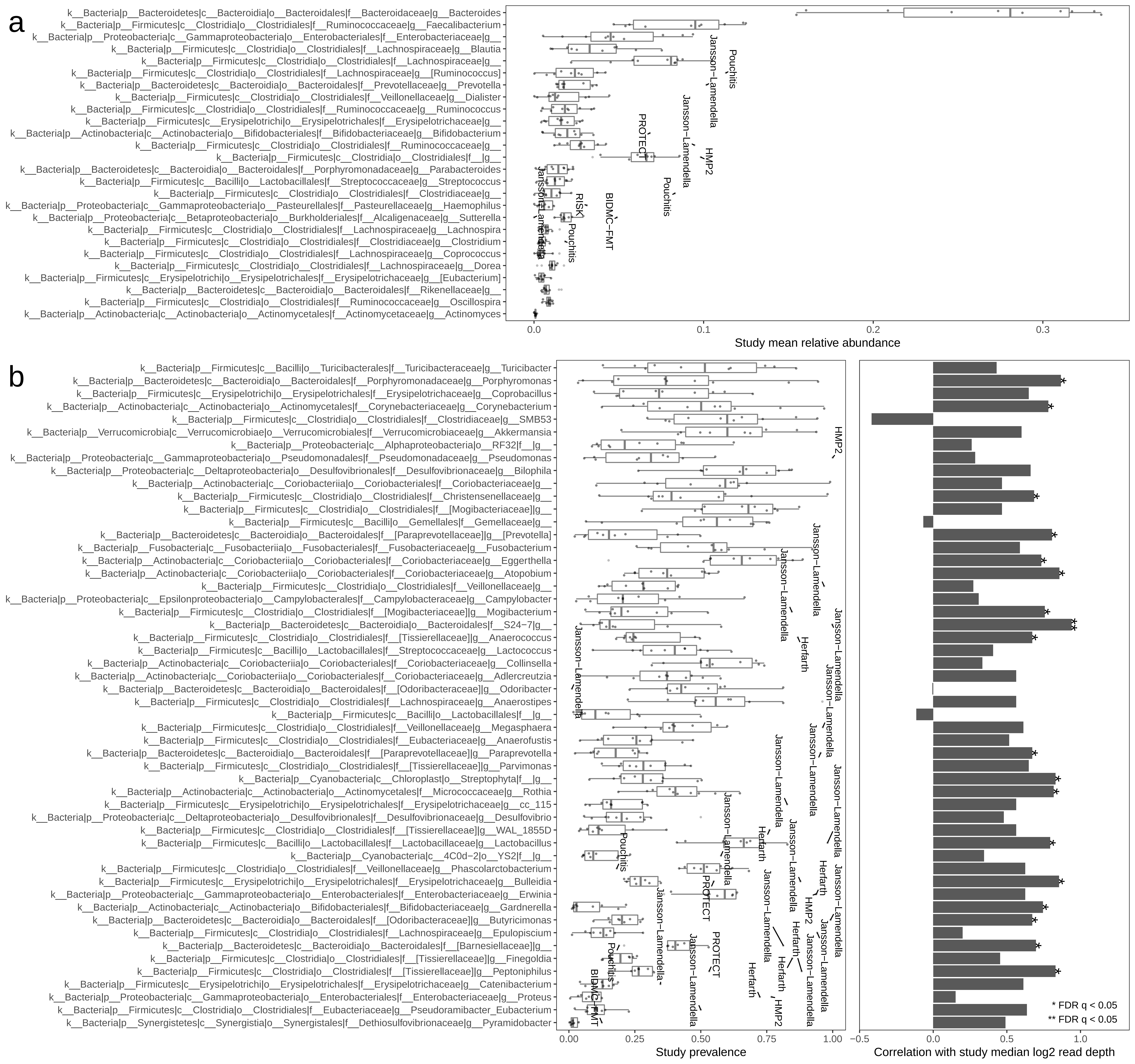

### suppFig3.pdf

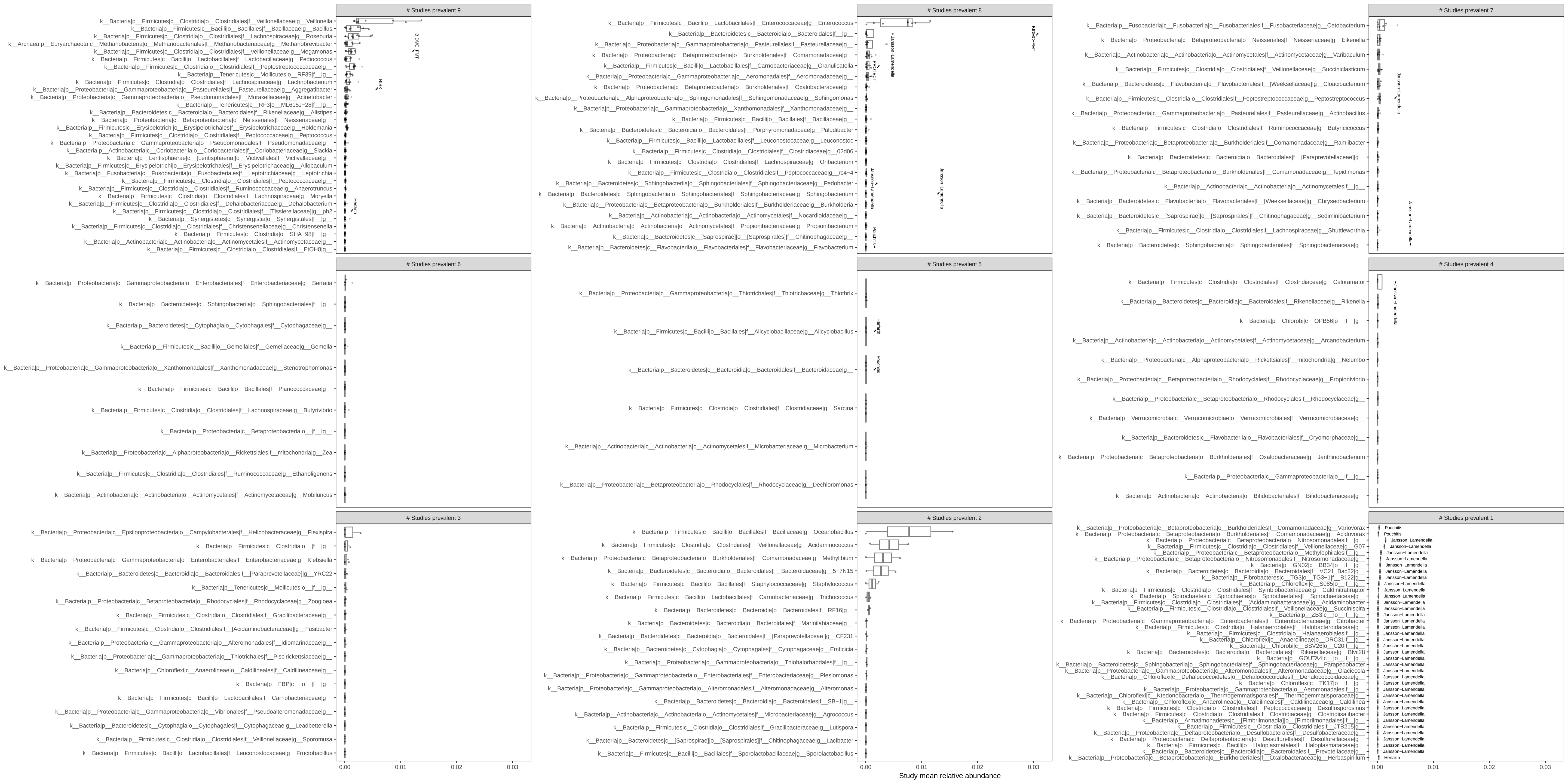

### suppFig4.pdf

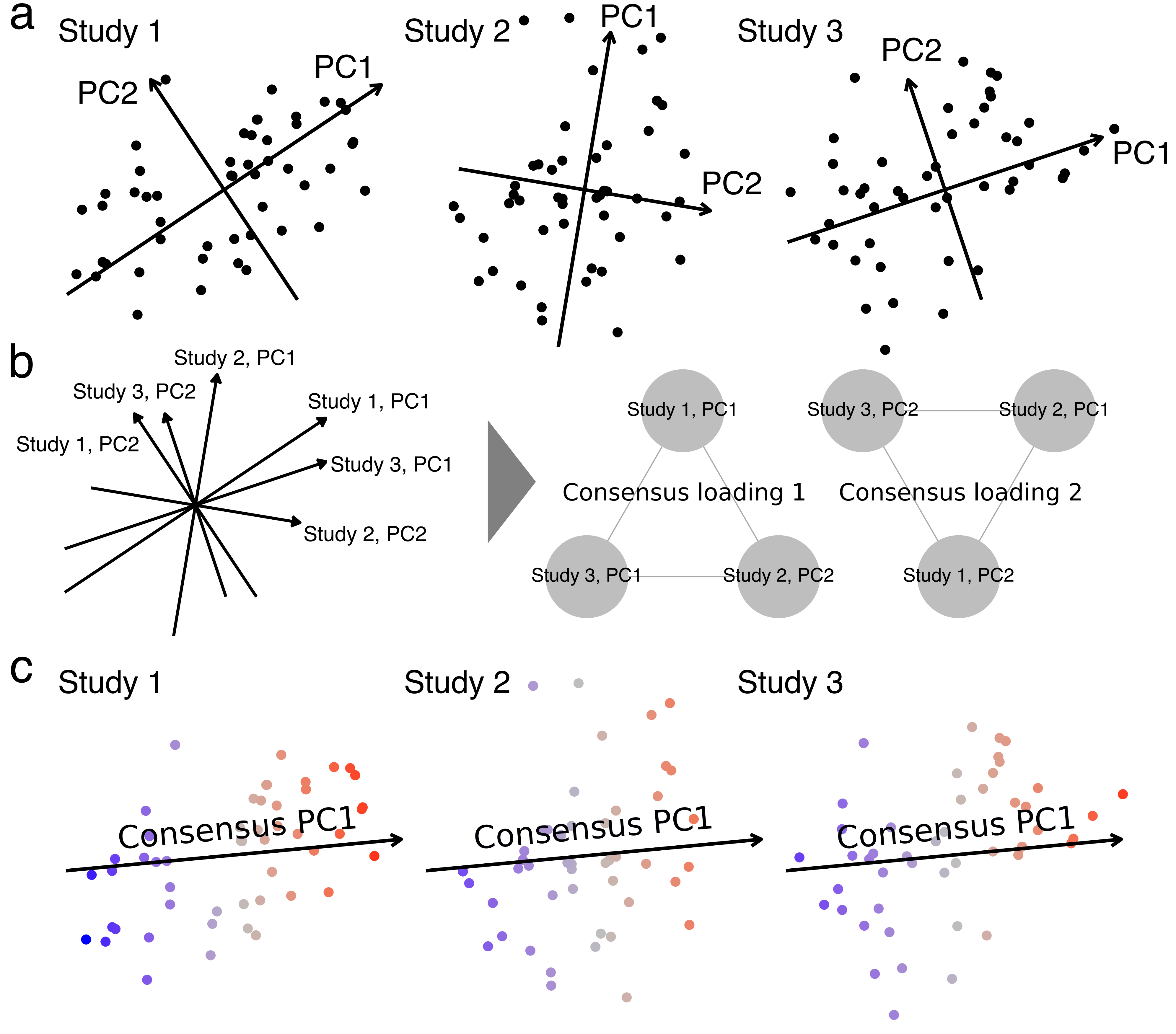

### suppFig5.pdf

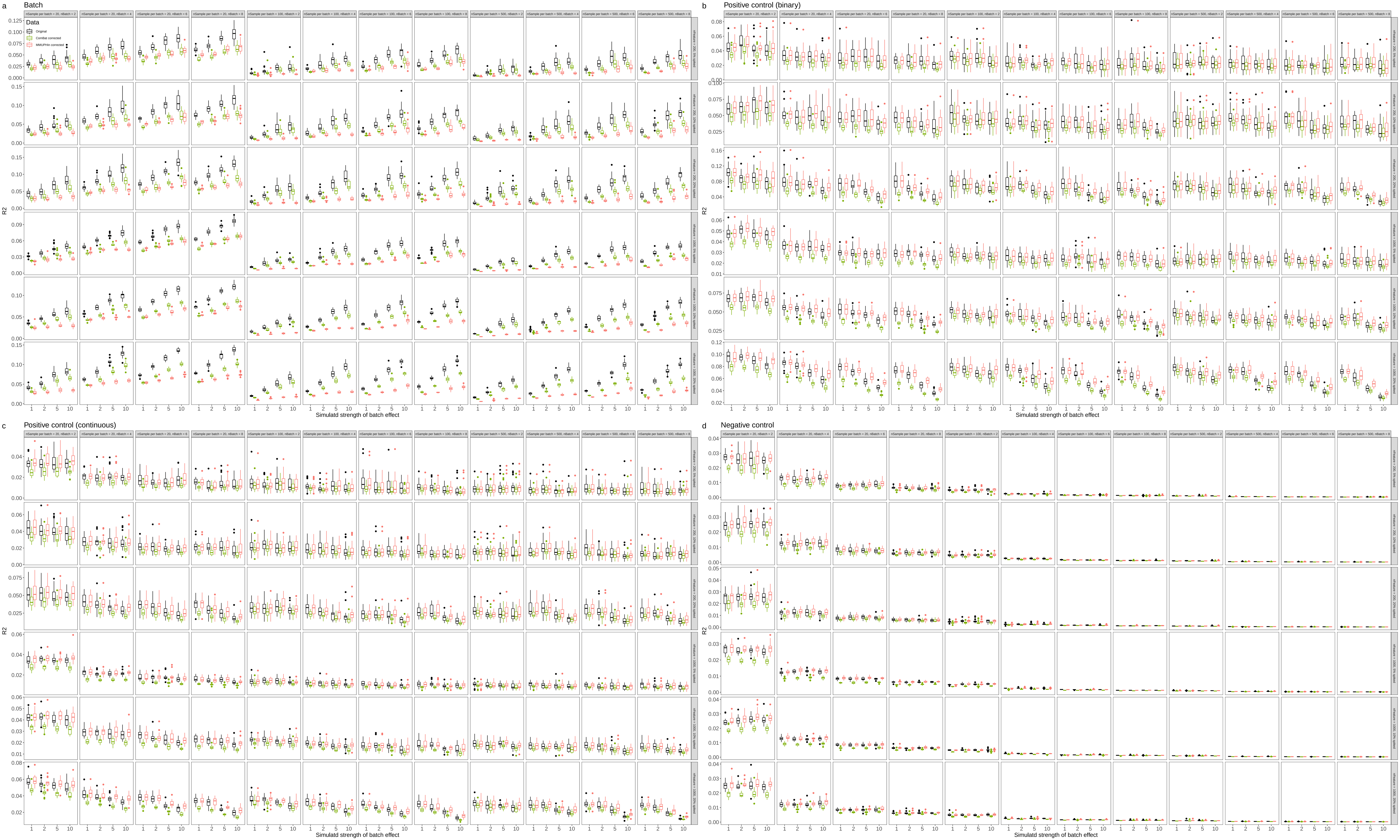

### suppFig6.pdf

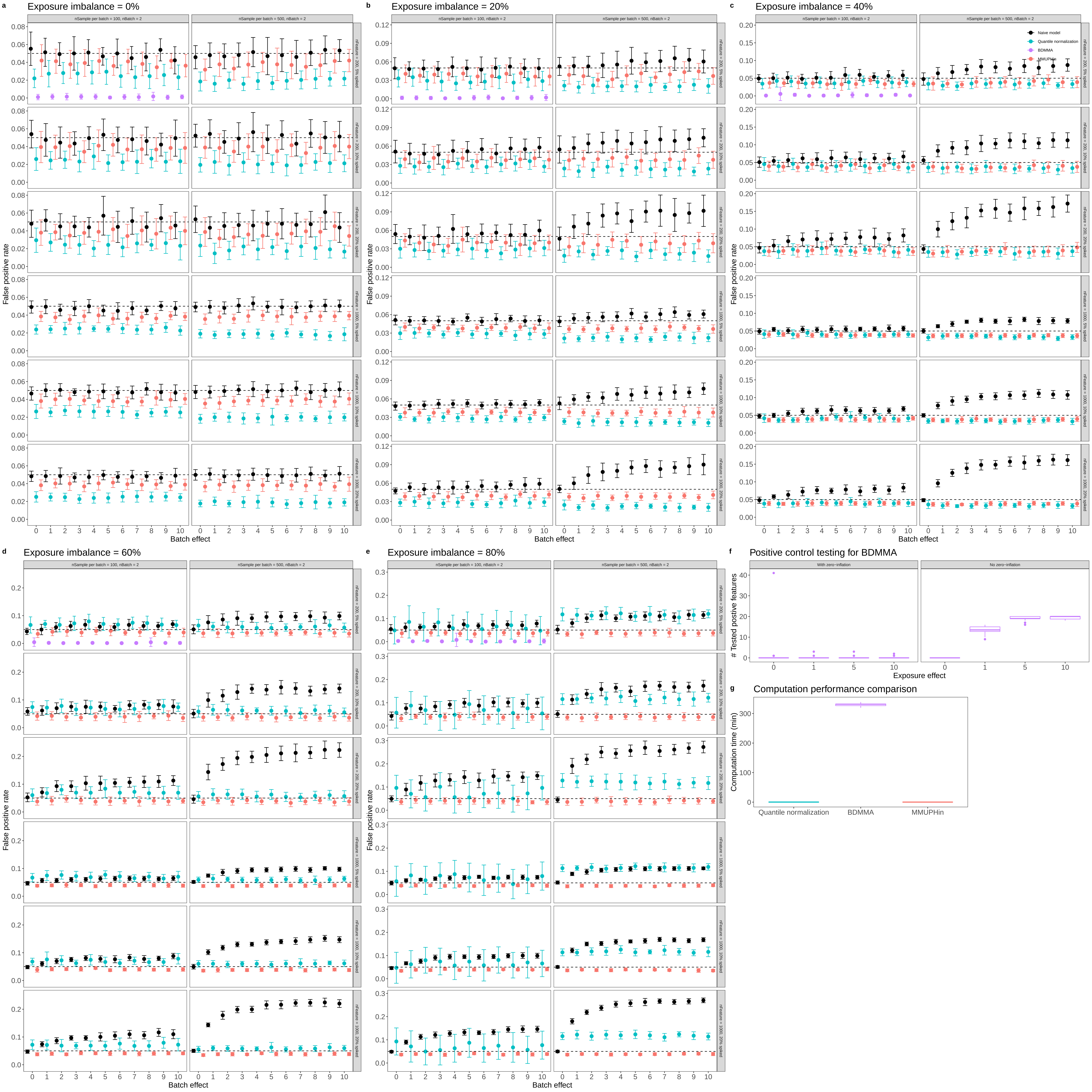

### suppFig7.pdf

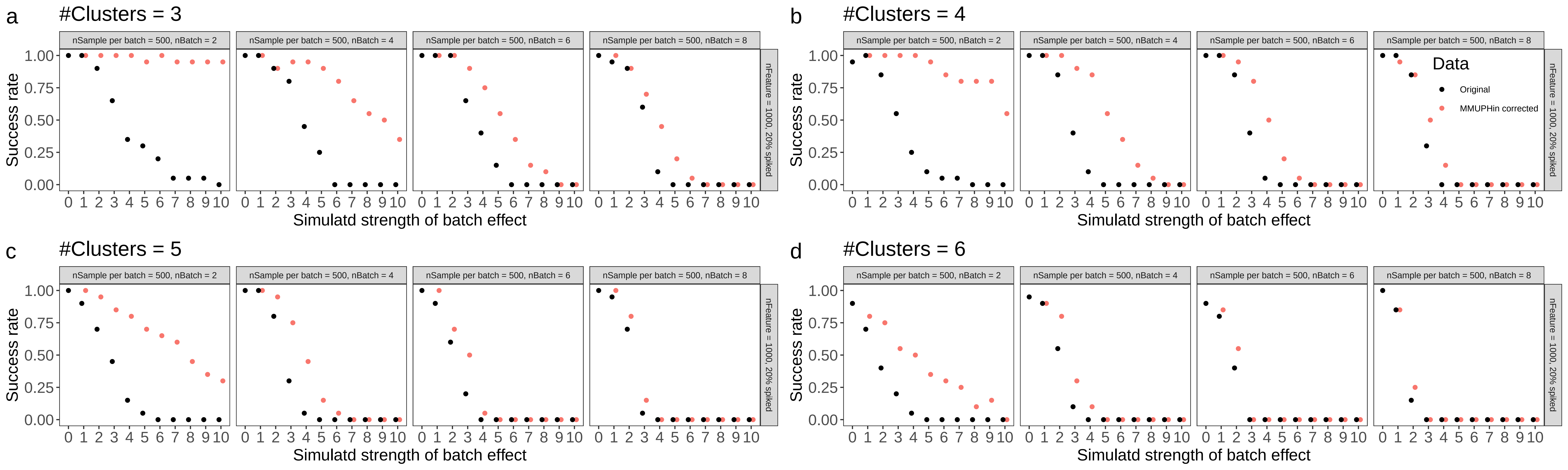

### suppFig8.pdf

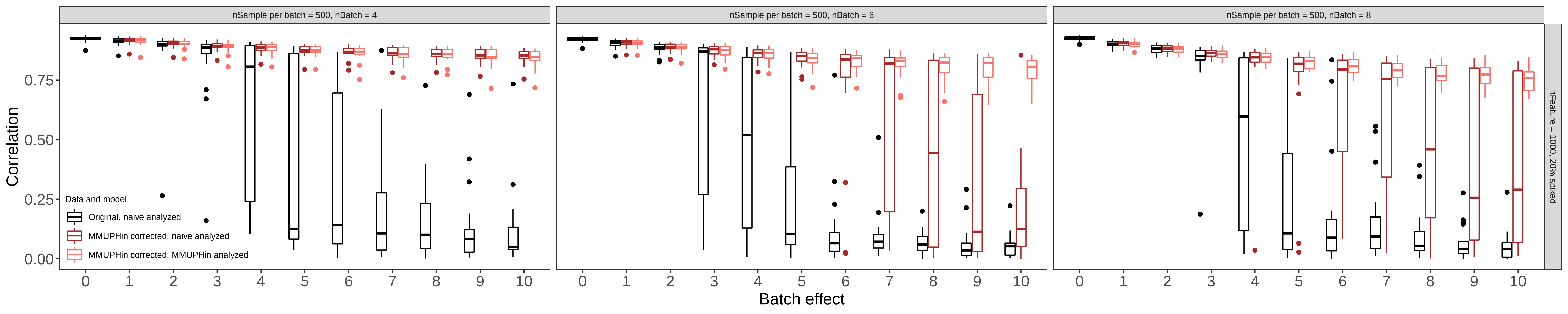

### suppFig9.pdf

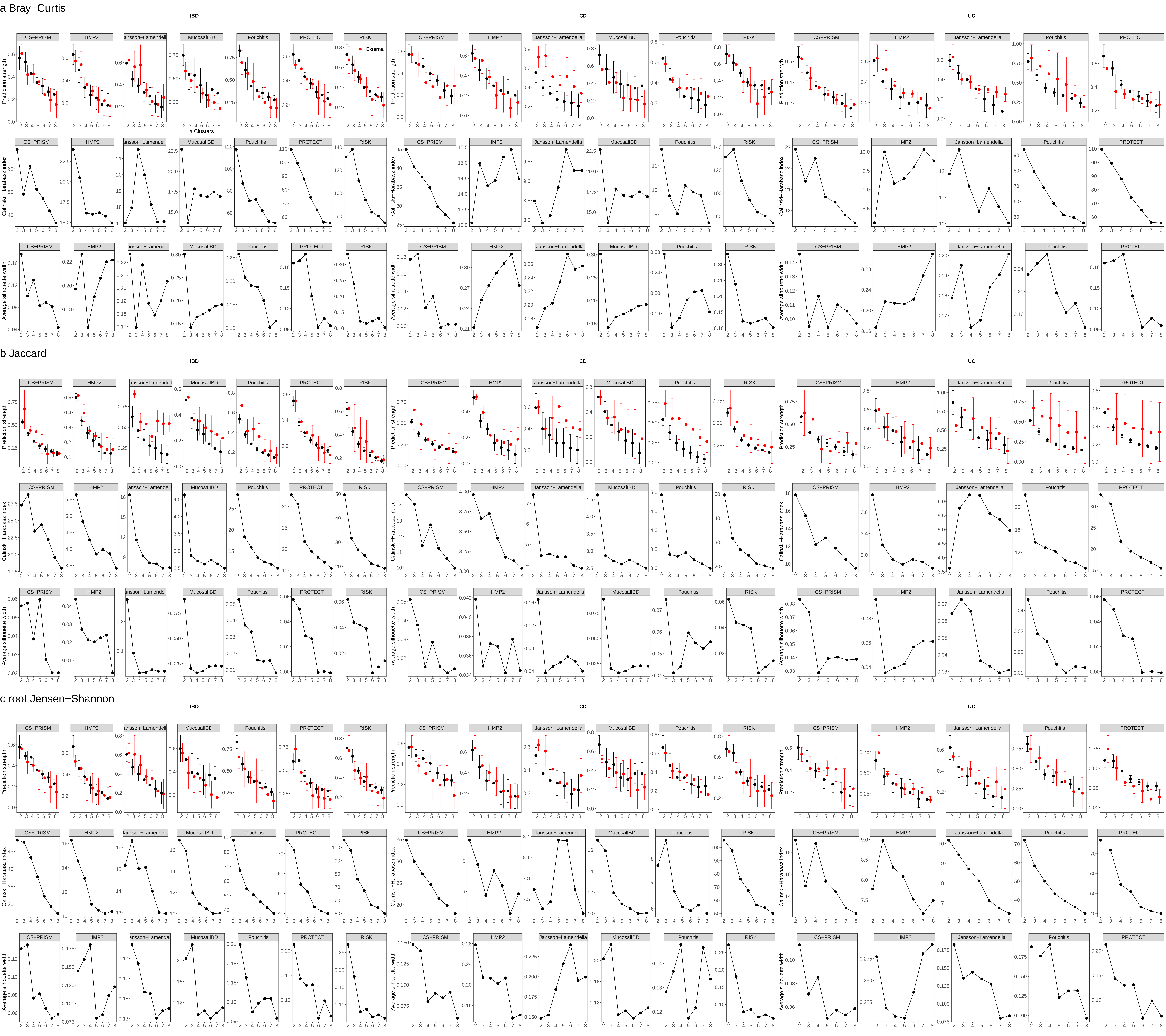

### suppFig10.pdf

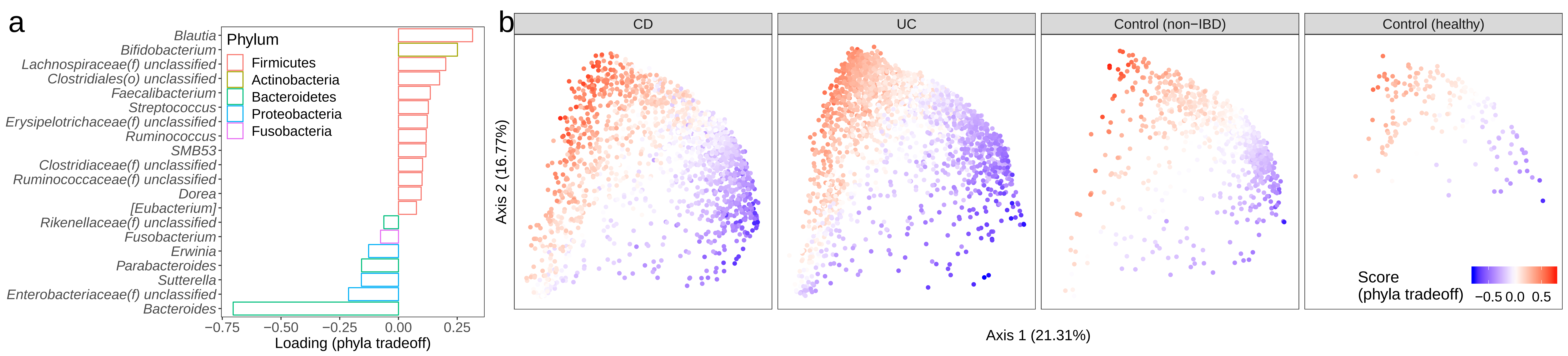

### suppFig11.pdf

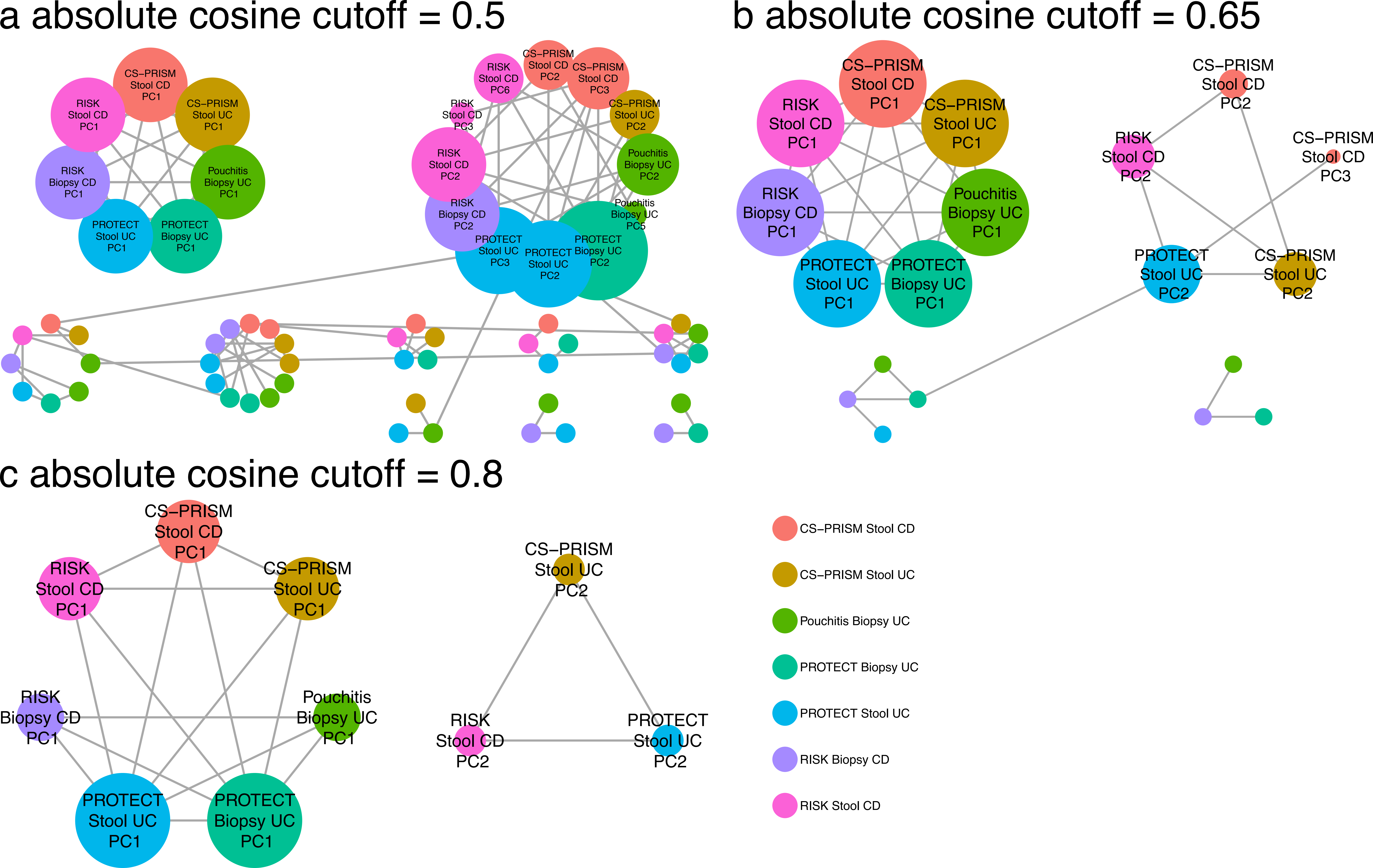

### suppFig12.pdf

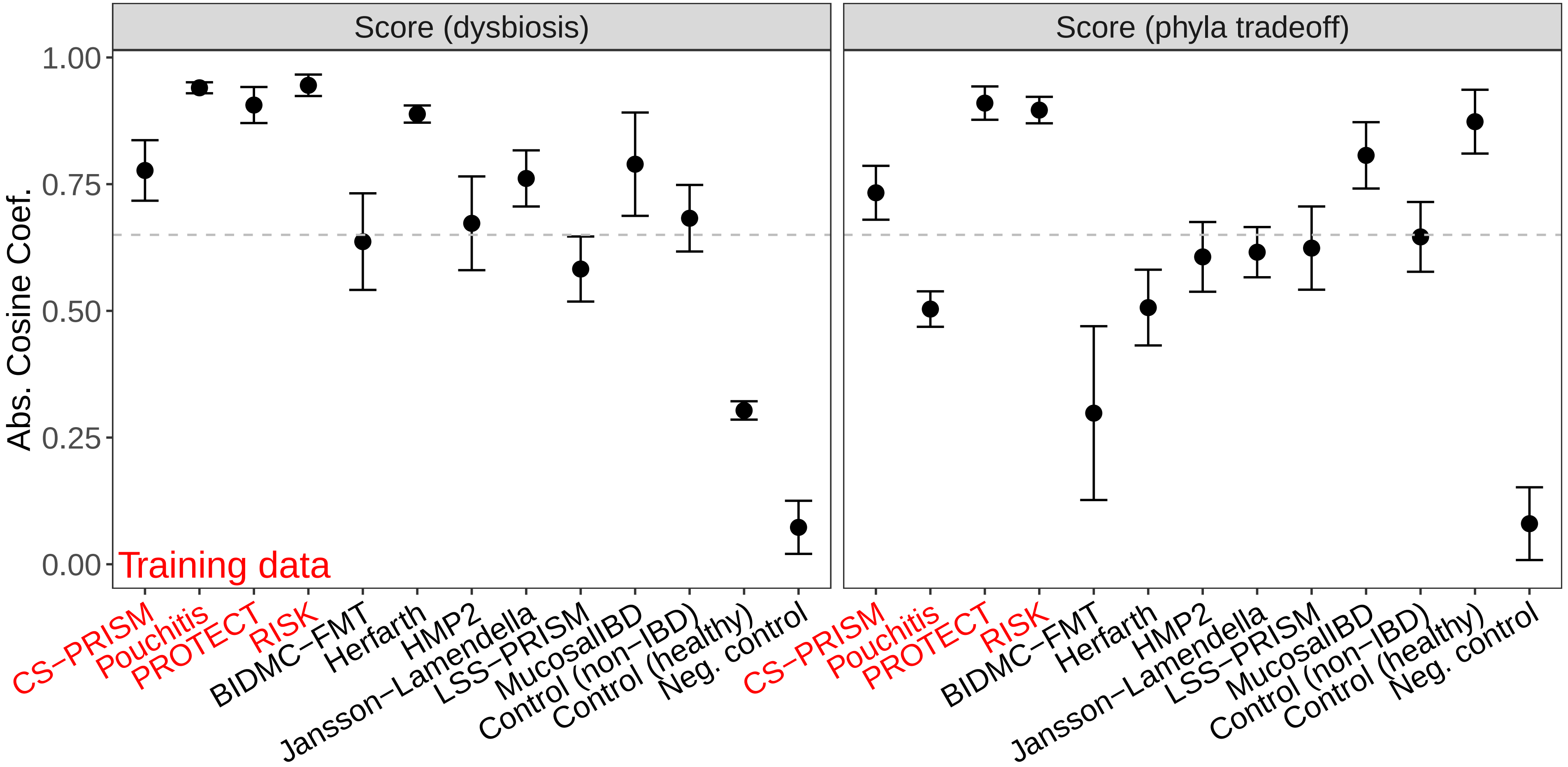

### suppFig13.pdf

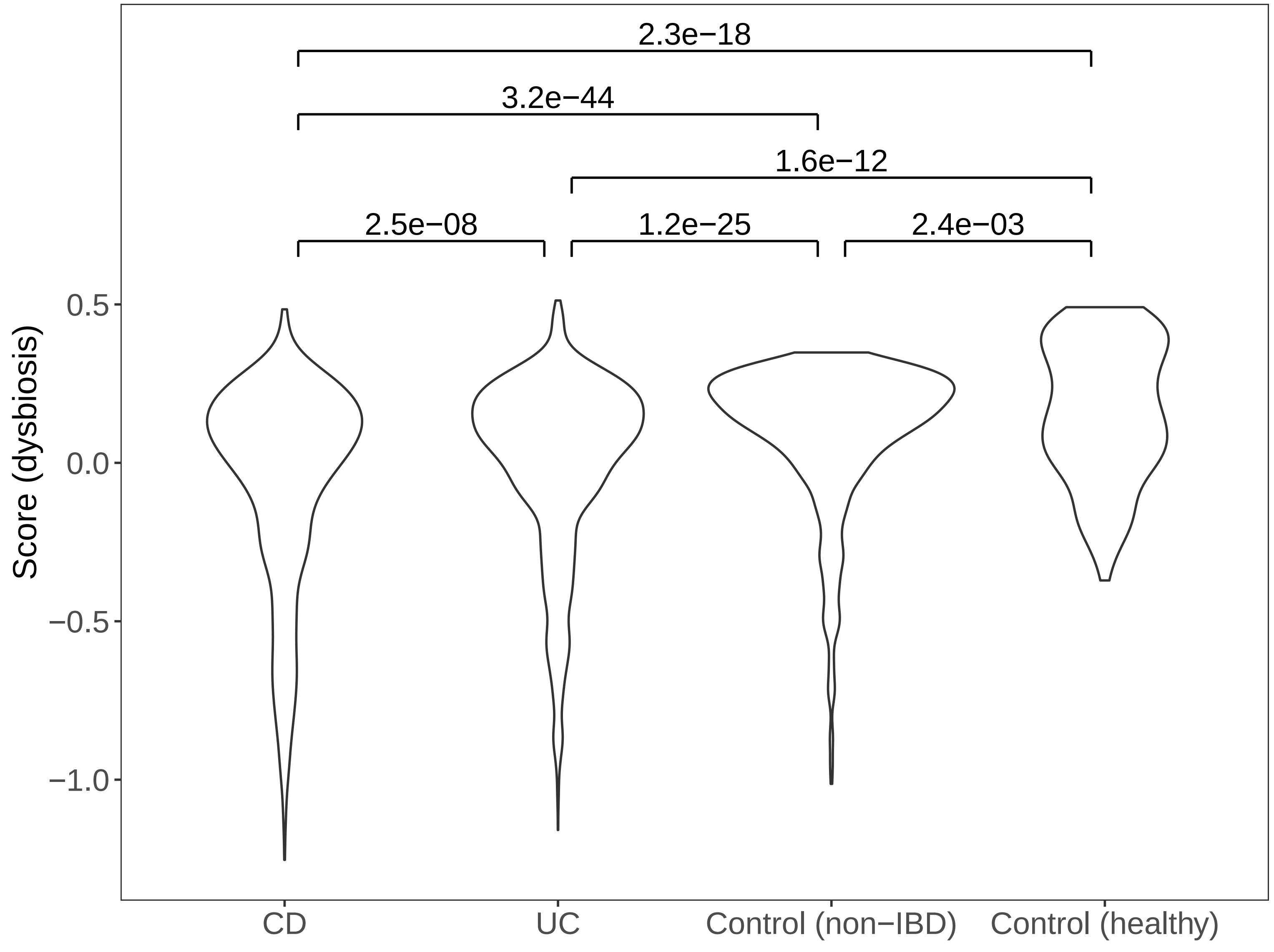
